# Kinematic and neuromuscular characterization of cognitive involvement in gait control in healthy young adults

**DOI:** 10.1101/2022.07.01.498421

**Authors:** Valentin Lana, Julien Frère, Vincent Cabibel, Tristan Réguème, Nicolas Lefèvre, Elodie Vlamynck, Leslie M. Decker

## Abstract

The signature of cognitive involvement in gait control has rarely been studied using both kinematic and neuromuscular features. The present study aimed to address this gap. Twenty-four healthy young adults walked on an instrumented treadmill in a virtual environment under two optic flow conditions: normal (NOF) and perturbed (POF, continuous mediolateral pseudorandom oscillations). Each condition was performed under single-task and dual-task conditions of increasing difficulty (1-, 2-, 3-back). Subjective mental workload (raw NASA-TLX), cognitive performance (mean reaction time and d-prime), kinematic (steadiness, variability and complexity in the mediolateral and anteroposterior directions) and neuromuscular (duration and variability of motor primitives) control of gait were assessed. The cognitive performance and the number and composition of motor modules were unaffected by simultaneous walking, regardless of the optic flow condition. Kinematic and neuromuscular variability was greater under POF compared to NOF conditions. Young adults sought to counteract POF by rapidly correcting task-relevant gait fluctuations. The depletion of cognitive resources through dual-tasking led to reduced kinematic and neuromuscular variability and this occurred to the same extent regardless of simultaneous working memory (WM) load. Increasing WM load led to a prioritization of gait control in the mediolateral direction over the anteroposterior direction. The impact of POF on kinematic variability (step velocity) was reduced when a cognitive task was performed simultaneously, but this phenomenon was no modulated by WM load. Collectively, these results shed important light on how young adults adjust the processes involved in goal-directed locomotion when exposed to varying levels of task and environmental constraints.

**NEW & NOTEWORTHY:** The kinematic and neuromuscular signatures of cognitive involvement in gait control have rarely been studied jointly. We sought to address this issue using gait perturbation and dual-task paradigms. The protocol consisted of a fixed-speed treadmill walk to which visual and cognitive constraints were applied separately and together. The results revealed that young adults optimally regulated their gait to cope with these constraints by maintaining relatively stable muscle synergies and flexibly allocating attentional resources.

## INTRODUCTION

Walking is a common but complex human behavior that relies on central pattern generators, which are regulated by supraspinal control and somatosensory feedback (1, 2). To navigate in the complex environments of everyday life requires the brain to continuously adapt to external constraints, such as obstacles or irregular surfaces (3) and to cognitively demanding internal tasks (4). To this end, the central nervous system controls locomotion in a hierarchical and distributed manner by modulating the activity of its control subsystems such as central pattern generators (2, 5, 6). These networks are then modulated by descending cortical and brainstem pathways, as well as by sensory feedback to regulate ongoing movement (2, 7). Then, control of walking is never strictly under the control of either automatic or executive control processes. There is a balance between the two processes, depending on the demands of the task and the capabilities of the individual (8). Typically, goal-directed walking requires executive control processes (9). Collectively, these mechanisms allow for the control of multiple joints and muscle groups in order to cope with the many associated large degrees of freedom (10).

At neuromuscular level, to simplify the development of complex movements, it has been hypothesized that the central nervous system (CNS) relies on a limited number of low-dimensional structures called muscle synergies (11, 12). In this sense, the observed changes reflect the flexibility of the modular organization. Each synergy contains a spatial component (or a motor module) that represents the involvement of each muscle within the synergy and a temporal component (or a motor primitive) that reflects its activation profile (13). On the one hand, the number and composition of motor modules is invariable under sensory perturbations in healthy young and older adults with no history of falls (14). On the other hand, Santuz et al. (15) observed an increase in the duration of motor primitives when walking was perturbed by continuous mediolateral oscillations of the support surface (*i.e.* mechanical perturbations). This increase was interpreted as a compensatory mechanism implemented by the CNS in response to increased postural threat (16). In line with Desrochers et al. (17), we propose that the organization of muscle synergies within the spinal cord simplifies the neural control of locomotion, freeing up resources for cortical structures to handle goal-directed actions, such as coping with gait perturbations or performing concomitant cognitive tasks. To our knowledge, no study has assessed the extent to which the modification of gait control under visual perturbations requires attentional resources and modulates underlying muscle synergies. Only Walsh (18) assessed the modulation of muscle synergies during walking and running when a concomitant cognitive task (counting backwards by 7s) was added. However, the author observed no significant effects of the dual task on motor primitive duration.

In order to better understand and characterize the cognitive involvement in gait control under visual perturbations and/or conditions of cognitive depletion, it is necessary to study the mechanisms of kinematic regulation in parallel with those of neuromuscular regulation. Dingwell & Cusumano (19) have proposed and demonstrated that Detrended Fluctuations Analysis can be used to detect the level of CNS intervention in the regulation of motor control. A tightly regulated variable implies active control to rapidly correct step-to-step deviations and suggests the intervention of executive functions (“executive control”, Clark, 8), whereas parameters that are not tightly regulated fluctuate freely and would result from a complex self-organizing system (20) (“automatic control”, Clark, 8). On a fixed-speed treadmill, these authors showed that step velocity was tightly regulated. Subsequently, another study observed a deterioration in step velocity regulation during walking as the attentional demand of the concomitant cognitive task increased (9). This confirms that executive functions (*e.g.* divided attention, working memory), which involve executive functions and coordinate goal-directed behavior (21, 22), are required to implement this regulation. This result remains to be confirmed in other contexts. Recently, Dingwell and Cusumano (23) have shown that, in the context of treadmill walking, in addition to step velocity, step width and, to a lesser extent, lateral body position (which corresponds to the distance between the center of the treadmill and the midpoint between the two feet), are tightly regulated to maintain mediolateral balance during walking. Furthermore, it has been experimentally confirmed that when walking under continuous mechanical or visual lateral perturbation, step width and lateral body position were more variable but also more tightly regulated (24–26). However, no link with executive functions has been demonstrated to date, although previous studies have suggested this under unperturbed walking conditions (9).

In fact, only two studies have evaluated the interaction effect of visual (*i.e.* optic flow) perturbations on gait under dual-task conditions (4, 27). The results of these studies suggest that, under these conditions, participants’ attentional resources would be redirected towards the performance of the concomitant cognitive task at the expense of the visual environment (4, 28). In turn, this phenomenon could diminish the influence of mediolateral optic flow perturbations on gait control. However, these two studies have several methodological limitations, including the use of (i) non-executive cognitive tasks (*i.e.* reaction time task and Go/No Go tasks, respectively) and (ii) less ecological (*i.e.* more simplistic and less realistic) optic flow perturbations than those generally used (25, 29). Thus, the involvement of executive functions in optic flow processing during gait control remains to be defined.

The present study aimed to characterize cognitive involvement in gait kinematic and neuromuscular control in healthy young adults. We hypothesized that (i) optic flow perturbations lead to greater kinematic (*i.e.* step parameters) and neuromuscular (*i.e.* motor primitives) variability, along with tighter regulation of task-relevant gait fluctuations (motor load manipulation; 30), (ii) depletion of cognitive resources through dual-tasking leads to the opposite effects (cognitive load manipulation; 31), and (iii) the impact of optic flow perturbations on gait is reduced during concomitant performance of a cognitive task, this phenomenon becoming more pronounced as cognitive load increases.

## MATERIALS AND METHODS

### Participants

Twenty-four healthy young adults (21.67 ± 2.28 years; 12 men and 12 women) took part in this study. Prior to the experiment, participants were interviewed regarding their health history. Exclusion criteria were: uncorrected visual impairment, any lower limb injury within the last six months, pain, functional limitations or any discomfort that may affect walking, history of vestibular, neurological or musculoskeletal disorders, body mass index of 30 or higher, sedentary lifestyle, and any medication altering the cognitive or physical abilities. The physical activity level was assessed by an adapted version of the question 1 of the Modifiable Activity Questionnaire (32). The lower limb laterality was determined by the leg used to kick a ball (33). All participants gave their written informed consent prior to the experiment. The study was approved by the French Research Ethics Committee CERSTAPS (IRB00012476-2020-30-11-76) and conducted in accordance with the principles of the Declaration of Helsinki regarding human experimentation.

### Experimental procedures

The experiment was conducted in the immersive room (dimensions: 4.80 × 9.60 × 3 m) of the Interdisciplinary Centre for Virtual Reality (CIREVE) at the University of Caen Normandie. The training session started with a familiarization of the participants to the 2 × 0.5 m walking treadmill (M-Gait, Motekforce Link, The Netherlands) followed by the determination of their preferred walking spe ed according to the method of Jordan et al. (34). Briefly, participants initially started at a relatively low speed (*e.g.* 0.6 m.s^−1^), which was gradually increased by 0.05 m.s^−1^ increments until they reported being at their preferred walking speed. Then, walking speed was further increased by approximately 0.42 m.s^−1^ and decreased in 0.05 m.s^−1^ decrements until the participants reported once again being at their preferred walking speed. The procedure was repeated until the ascending and descending speeds reported by the participants were close (difference below 0.1 m.s^−1^). Afterwards, the participants were familiarized with the auditory N-back task with three levels of WM load (*i.e.* 1-back, 2-back, and 3-back, *see task description below*). Each familiarization trial lasted at least 1 minute.

The testing session was composed of three blocks performed in a randomized order: (1) three N-back conditions in a seated position (*i.e.* single-task cognitive performance), (2) walking under normal optic flow (NOF) conditions, and (3) walking under perturbed (*i.e.* continuous mediolateral pseudo-random oscillations) optic flow conditions (POF). In the last two blocks, walking tasks were performed under both single-task walking (STW) and dual-task walking (DTW) conditions (*i.e.* responding to the N-back tasks). Participants were instructed to walk naturally while looking straight ahead. The treadmill speed was adjusted to their preferred walking speed. The blocks (2) and (3) began and ended with a STW condition while the three DTW conditions were performed in a randomized order between the two STW conditions (35). Under dual-task conditions, no task priority instructions were given (35). Each condition lasted 3 minutes. Hence, a total of thirteen experimental conditions were performed (Figure 1).

**Figure 1.**
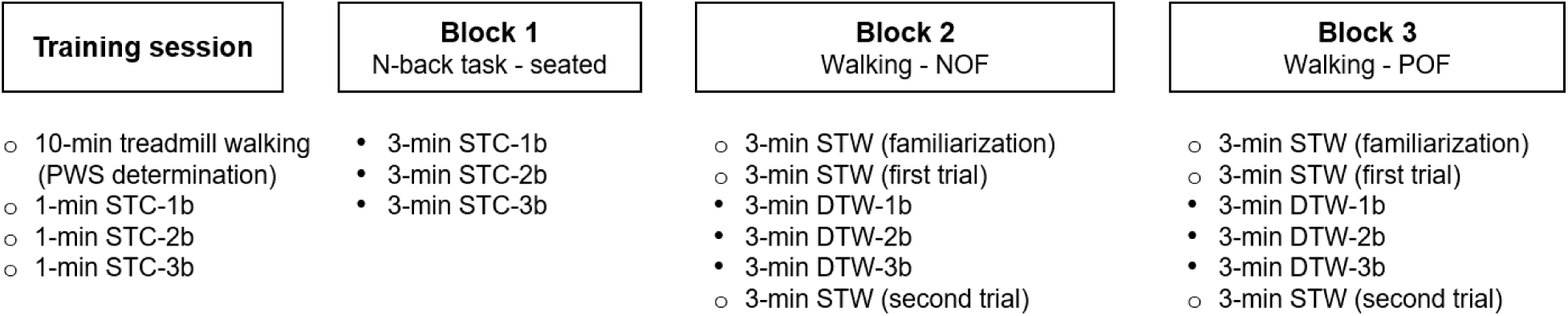
Experimental procedures. A training session was first performed, consisting of: 1) a 10 - minute familiarization period with treadmill walking, 2) the determination of the participant’s preferred walking speed on the treadmill using the method of Jordan et al. (2007), and 3) a 1-minute familiarization trial with each of the three auditory N-back tasks, i.e. 1-back (1b), 2-back (2b), and 3-back (3b). Next, three experimental blocks were performed in a randomized order: 1) N-back tasks in a seated position (STC: single-task cognitive performance), 2) walking under normal optic flow (NOF), 3) walking under perturbed optic flow (POF, continuous mediolateral pseudo-random oscillations). In the last two blocks, participants began and ended with a single-task walking (STW) condition (i.e. free walking), while the three dual-task walking (DTW) conditions were performed in a randomized order between the two STW conditions. The experimental conditions lasted 3 minutes. Empty circles correspond to familiarization trials while solid circles correspond to experimental conditions.

At the end of each condition, participants were asked to complete the raw NASA task load index (NASA-TLX), a subjective multidimensional assessment questionnaire of perceived workload, on a digital tablet (36, 37). In this questionnaire, three subscales relate to the demands imposed on the participant (*i.e.* physical, mental and temporal demands), and three others to the interaction of the participant with the task (*i.e.* effort, frustration, performance).

#### Manipulation of optic flow

Participants were asked to walk in a virtual environment composed of a street in ancient Rome in which white statues of 2.4 m height were spaced every 3 meters to increase motion parallax (37, 38, Figure 2).

**Figure 2.**
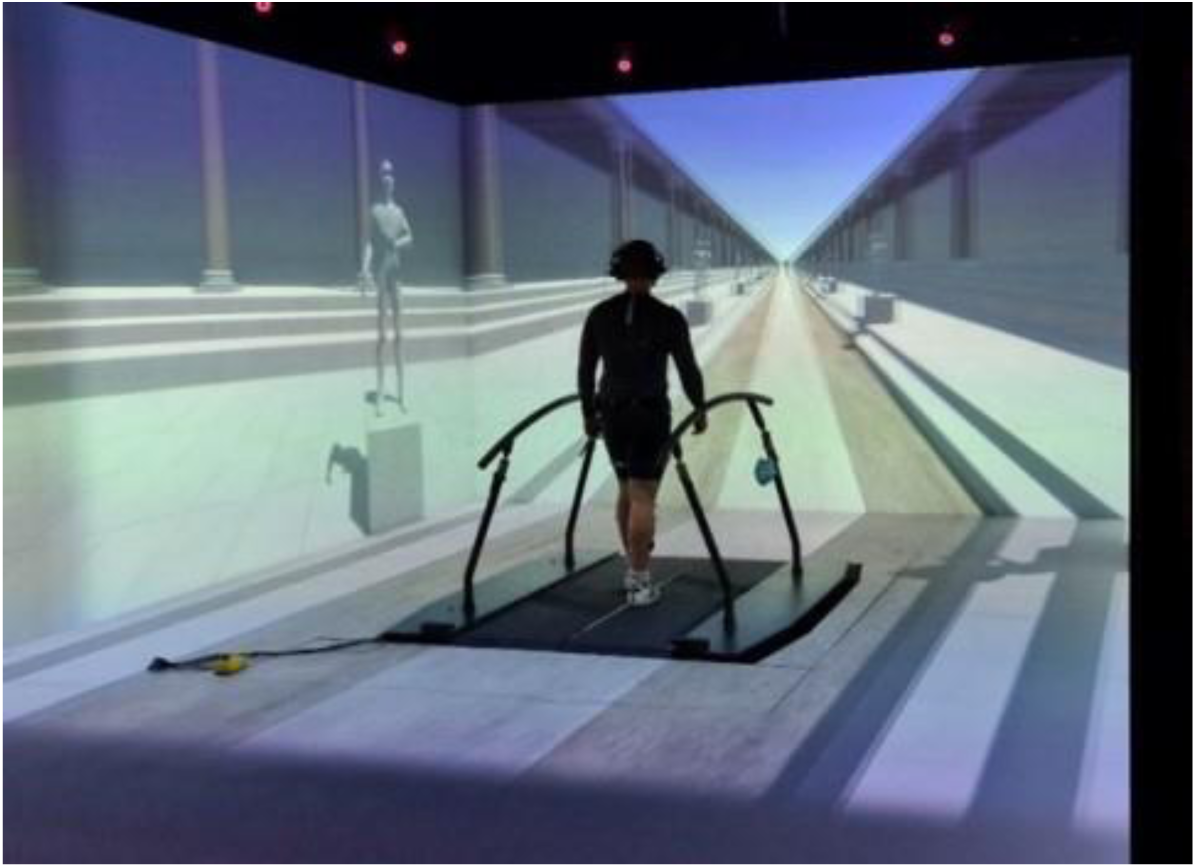
Experimental setup. Photograph of a participant walking on the M-Gait dual-belt instrumented treadmill (Motekforce Link, Amsterdam, The Netherlands) in a virtual environment that was developed specifically for the study. The participant wears a suit with 27 retroreflective markers (4-mm diameter) placed on specific anatomical landmarks according to an adaptation of the Plug-in-Gait full-body model (Vicon Motion Systems Ltd., Oxford, UK).

Under all conditions, the speed of the optic flow in the anteroposterior direction corresponded to the speed of the treadmill belts. Under perturbed optic flow conditions, the visual field oscillated in the mediolateral direction around the center line of the treadmill. The perturbation consisted of a pseudo-random sum of four sinusoids with an amplitude of 0.25 m, already used in the literature (25, 29):

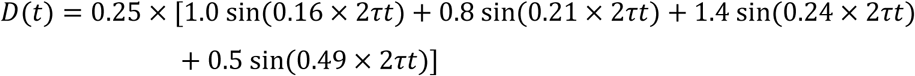

where D(t) is the lateral translation distance (m).

#### Manipulation of working memory load

The N-back task was used in this study. This task consists of a continuous sequence of auditory stimuli, presented one by one, in which the participant must determine whether the stimulus currently presented is the same as the stimulus presented N trials previously. Thus, the ‘N’ factor (*i.e.* number of items stored in WM) increases the task difficulty and hence the WM load (40). Moreover, it involves both working memory and information processing, which has a greater impact on gait than tasks involving inhibition or visuospatial cognition, particularly during perturbed walking (41). In the present study, WM load was parametrically manipulated across three levels of an auditory N-back task (*i.e.* 1-back, 2-back, 3-back). 2-back and 3-back tasks tax executive functions while 1-back task involves few or no executive functions and are more akin to attentional tasks (42, 43). According to Bayot et al. (44), executive functions “encompass the higher cognitive processes involved in the cognitive control of non-routine, goal-directed behaviors”. The auditory modality was chosen to avoid interfering with optic flow perturbations.

Using noise-cancelling headphones, participants heard the letters “E - A - I - O - R - T” pronounced easily and clearly in French, and were asked to answer “Yes” when the letter heard was the same as that presented N trials earlier (*i.e.* 1-back, 2-back, 3-back). Responses were recorded using a microphone. The letters were presented with an inter-stimulus interval of 1800 to 2200 ms (2000 ± 10%) to avoid the influence of rhythm on walking (35). Each letter was presented for a duration of 500 ms. Each sequence consisted of one-third targets (*i.e.* 90 stimuli for which the participant had to answer “Yes“; 41), along with 9 (10%), 13 (14.4%) and 16 (17.8%) lures for the 1-back, 2-back and 3-back, respectively (46). Each condition lasted 3 minutes.

#### Kinematic recording

The participants were equipped with 4-mm diameter retroreflective markers, fixed to specific anatomical landmarks on the body, on top of fitted clothing (47). The model used for marker placement was an adaption of the Plug-in Gait full-body model (48): only the head, trunk and lower limb markers were used for a total of 27 markers. The three-dimensional (3D) marker positions were collected at 200 Hz using 12 infrared cameras (Vero 2.2, Vicon Motion Systems Ltd., Oxford, United Kingdom). The origin of the reference frame was at the treadmill’s center. Any gap in the raw motion capture data were filled using Vicon Nexus software (version 2.8.1, Oxford Metrics, Oxford, United Kingdom). The 3D marker trajectories were then imported into MATLAB® R2020a (The MathWorks Inc., Natick, Massachusetts, USA) for further analyses.

#### Electromyographic recording

Once the skin prepared (*i.e.* shaving, light abrasion, cleaning with alcohol solution), surface electromyographic electrodes (EMG Trigno Snap Lead Sensors, Delsys, Inc., Natick, Massachusetts, USA) were placed on the muscles: tibialis anterior (TA), soleus (SOL), medial gastrocnemius (GAS), vastus medialis (VM), rectus femoris (RF), semitendinosus (ST), biceps femoris (BF), gluteus medius (Gmed) of the dominant lower limb (conversion A/D, 1000 Hz, 10 V). Electrodes were placed longitudinally with respect to the arrangement of muscle fibers (49) and located according to the recommendations from Surface EMG for Non-Invasive Assessment of Muscles (SENIAM; 50).

### Data analysis

#### Subjective mental workload

In all conditions, subjective mental workload was estimated based on the raw NASA task load index (NASA-TLX, with a maximum possible score of 100 points), that is the sum of the scores of the six subscales (51).

#### Cognitive task performance

Mean reaction time for target trials (RT, seconds) and d-prime (d’, a.u.) were used to assess performance in N-back tasks (52–54). RT was measured only for correct responses and corresponded to the time elapsed between the onset of the auditory stimulus and the start of the participant’s response. The d’ was the difference between the Z-score obtained from hit rate, Z_Hit_, and that obtained from the false alarm rate (FA, *i.e.* participant answers “Yes” when there was no target), Z_FA_ (d’ = Z_Hit_ - Z_FA_; 51). The higher the hit rate and the lower the FA, the better the participant’s ability to discriminate target and non-target letters. In the present experiment, a d’ of 4.5 indicated 100% correct responses and no false alarms.

#### Gait kinematics

The kinematic data were low-pass filtered using a fourth-order zero-phase Butterworth filter with a cut-off frequency of 6 Hz. Heel strikes were computed from the heel marker trajectory using the algorithm of Zeni et al. (56). Next, step velocity, step width and lateral body position were extracted, as previously done (23, 25). The mean (steadiness), standard deviation (variability) and statistical persistence/anti-persistence of the time series (complexity) of these variables were then calculated. To quantify how fluctuations were regulated from one step to the next, the Detrended Fluctuations Analysis scaling exponent (α), which assesses the presence and strength of statistical persistence or anti-persistence in a given time series, was used (57–60). An α < 0.5 suggests that the time series contains anti-persistent correlations, *i.e.* subsequent deviations are more likely to be in the opposite direction, consistent with a tightly controlled process. An α = 0.5 indicates uncorrelated deviations that are attributed to noise. An α ≈ 0.5 is typically exhibited by variables that are more tightly regulated. An α >> 0.5 (*i.e.* strong persistence) is typically exhibited by variables that are less tightly regulated (23, 25, 60). Given the length of our time series (290 steps), the scaling exponent (α) was computed following the recommendations of Phinyomark et al. (61). We also performed a direct control analysis of step-to-step fluctuations to complete the Detrended Fluctuations Analysis (62). For *q* ∈ {*V*, *W*, *z*_*B*_}, participants trying to maintain a constant average value, *q̅*, on the treadmill exhibit deviations away from *q̅* on any given step, 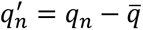. Each such deviation is corrected on the subsequent step with a corresponding change in the opposite direction, Δ*q*_*n*+1_ = *q*_*n*+1_ − *q*_*n*_. From plots of 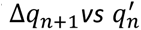, we computed (using least-squares regression) the linear slopes and strength of correlation (R²). Tightly controlled variables would be corrected quickly, yielding a slope value (M) close to −1 and a high R² value (23, 25). These variables, in particular those derived from the Detrended Fluctuations Analysis, have demonstrated their ability to detect the involvement of executive control processes in gait regulation: the more tightly these variables are regulated, the greater the involvement of executive control (9, 60, 63).

#### Gait electromyography

The raw EMG signals were band-pass filtered using a fourth-order zero-phase IIR Butterworth filter with cut-off frequencies of 50 and 450 Hz, then rectified and smoothed using a fourth-order low-pass zero-phase IIR Butterworth filter with a cut-off frequency of 10 Hz (16). After subtracting the minimum, the muscle activation profiles were normalized in amplitude to the average peak activation of the selected gait cycles (*i.e.* N = 80). Lastly, each gait cycle was time-normalized to 200 points by a spline interpolation. We excluded data from 9 participants due to poor EMG signal quality (*i.e.* movement-related artefacts). Therefore, EMG data from 15 right-handed participants were retained (21,67 ± 2,53 years old; BMI: 21,44 ± 1,90 kg.m^2^ ; 8 men and 7 women).

For each subject and each condition, previously processed EMGs were combined into a *m* × *t* matrix where *m* indicates the number of muscles and *t* their activation duration (original EMG matrix, EMGo). Next, muscle synergy extraction was performed by non-negative matrix factorization (NNMF; 59) of the *m* × *t* matrix corresponding to all gait cycles. The number of synergies *n* is specified a priori and the NNMF algorithm finds the properties of the synergies by filling in two matrices: a matrix *W* of dimensions *m* × *n*, which specifies the relative weighting of a muscle in each motor module, and a matrix *C* of dimensions *n* × *t*, which specifies the activation over time of each motor primitive for all gait cycles. The reconstructed EMG (EMGr) of a muscle is the sum of the contributions of all muscle synergies (65). To determine the minimum number of synergies required to correctly reconstruct EMGo, we quantified the Variance Accounted For (VAF; 66) as the ratio between the sum of squared error values and the sum of squared EMGo values 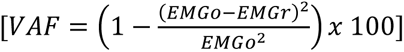. The number of synergies used corresponds to the one where VAF is greater than 90% with an increment of less than 5% for s+1 synergies (66). This approach is conservative and ensures high concordance between original and reconstructed EMG signals (65).

In each condition, the inter-cycle variability (67, 68) and duration (16, 69) of motor primitives were assessed using the variance ratio and the full width at half maximum (FWHM), respectively. The FWHM determines the number of points (percentage of the cycle) exceeding half of the maximum activation of each gait cycle. Next, the values obtained from all gait cycles were averaged to obtain a single FWHM value for each condition (15). The variance ratio corresponds to the variance at each cycle divided by the total variance. The variance ratio is determined for each motor primitive according to the following equation:

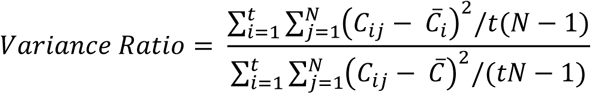

where *t* is the number of time points (*i.e.* 200 per cycle), *N* is the number of gait cycles over which variance ratio is evaluated, *C*_*ij*_ is the value of the *j*^*th*^ primitive waveform at time epoch *i*, *C̅*_*i*_ is the average primitive waveform at the time epoch *i* over *j* gait cycles, and *C̅* is the grand mean average primitive waveform (68).

### Statistical analysis

Linear mixed models (including a random intercept for each participant) were used to compare the effect of *WM load* (1-back, 2-back, 3-back) and *Cognitive condition* (single-task cognitive performance, dual-task cognitive performance under NOF, dual-task cognitive performance under POF) on cognitive performance (*i.e.* d’ and RT) and subjective mental workload (*i.e.* raw NASA-TLX score) dependent variables. Where main effects were significant, relevant pairwise comparisons were made using marginal means estimated with Tukey’s adjustment.

Performance during the second single-task walking (STW) conditions (*i.e.* performed at the end of the walking blocks) was not included in the analysis after observing and statistically confirming a disengagement or relaxation of the participants during these conditions (Fig. S1, *Supplementary materials*). Linear mixed models (including a random intercept for each participant) were used to compare the effect of *Optic flow* (NOF, POF) and *Walking condition* (STW, DTW-1b, DTW-2b, DTW-3b) on all kinematic dependent variables (*i.e.* mean, standard deviation, α exponent, slope and R² for step velocity, step width and lateral body position). Where main effects were significant, relevant pairwise comparisons were made using marginal means estimated with Tukey’s adjustment.

Beforehand, a chi² test was performed to assess the effect of *Optic flow* (NOF, POF) on the number of motor modules. A dot product (r) was used to evaluate the effect of *Walking condition* (STW, DTW-1b, DTW-2b, DTW-3b) on motor module composition. A threshold of r = 0.80 was defined below which motor modules would not be considered similar. Linear mixed models similar to those used for neuromuscular data were used to assess the effects of *Optic flow* (NOF, POF) and *Walking condition* (STW, DTW-1b, DTW-2b, DTW-3b) on motor primitives.

Linear mixed-models were fitted using the lme4 package (70) and tested on the basis of likelihood ratio tests (LRT) using the lmerTest package (71). For all variables, given that residuals are often not normally distributed (25), which was verified in our samples, model parameters were estimated using the pbkrtest package by computing 5000 parametric bootstraps (72). Post hoc comparison tests were performed using the emmeans package (v1.7.5). The significance threshold was set at 0.05. Effect sizes were also examined and reported for post hoc comparison tests using Cohen’s d (d < 0.2, 0.5 < d < 0.8 and d > 0.8, indicating weak, moderate and large effects, respectively). All statistical analyses were performed using R 4.0.5 (73).

## RESULTS

### Cognitive performance and subjective mental workload

A significant main effect of *WM load* was found for both d’ and RT (Table 1, *Table S1 for descriptive data*). Post hoc tests revealed a decrease in d’ (Figure 3A) and an increase in RT (Figure 3B) with increasing load. Precisely, for both dependent variables, 3-back performance was lower than 1-back and 2-back performances, and 2-back performance was also lower than 1-back performance. Significant main effects of *WM load* and *Cognitive condition* were found for the raw NASA-TLX score. Post hoc tests revealed an increase in subjective mental workload i) with increasing WM load, and ii) during dual-task cognitive performance under POF compared with single-task cognitive performance and dual-task cognitive performance under NOF. Thus, the resolution of a POF-induced sensory conflict during dual-task treadmill walking generated a higher subjective mental workload than in the absence of conflict. No interaction effect of *Cognitive condition* x *WM load* was observed (Figure 3C, Table 1). Therefore, the effect of WM load on subjective mental workload was no greater during dual-task cognitive performance, either in the presence or absence of POF, than during single-task cognitive performance.

**Table 1.**
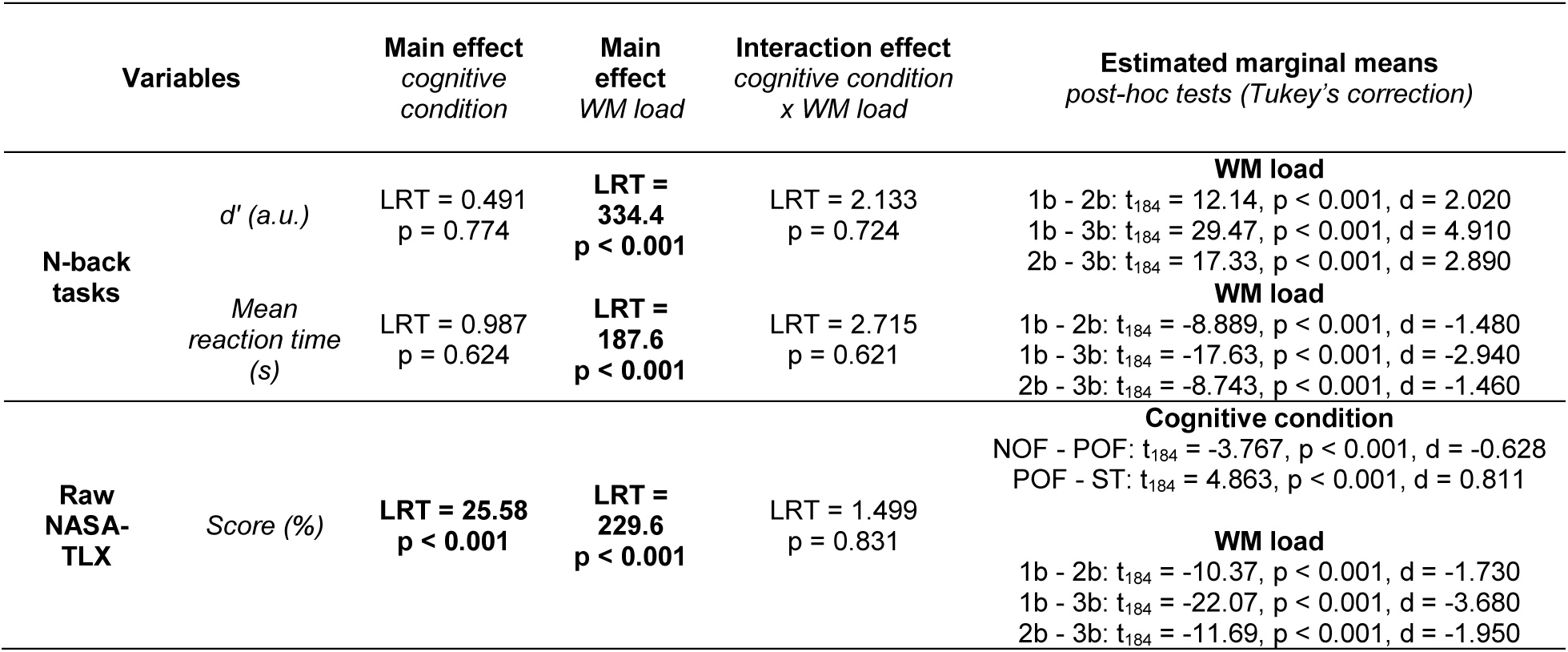
Statistical results for cognitive dependent variables: cognitive task performance, *i.e.* mean reaction time for target trials (RT, second) and d-prime (d’, a.u.), and subjective mental workload, *i.e.* raw NASA-TLX score.

**Figure 3.**
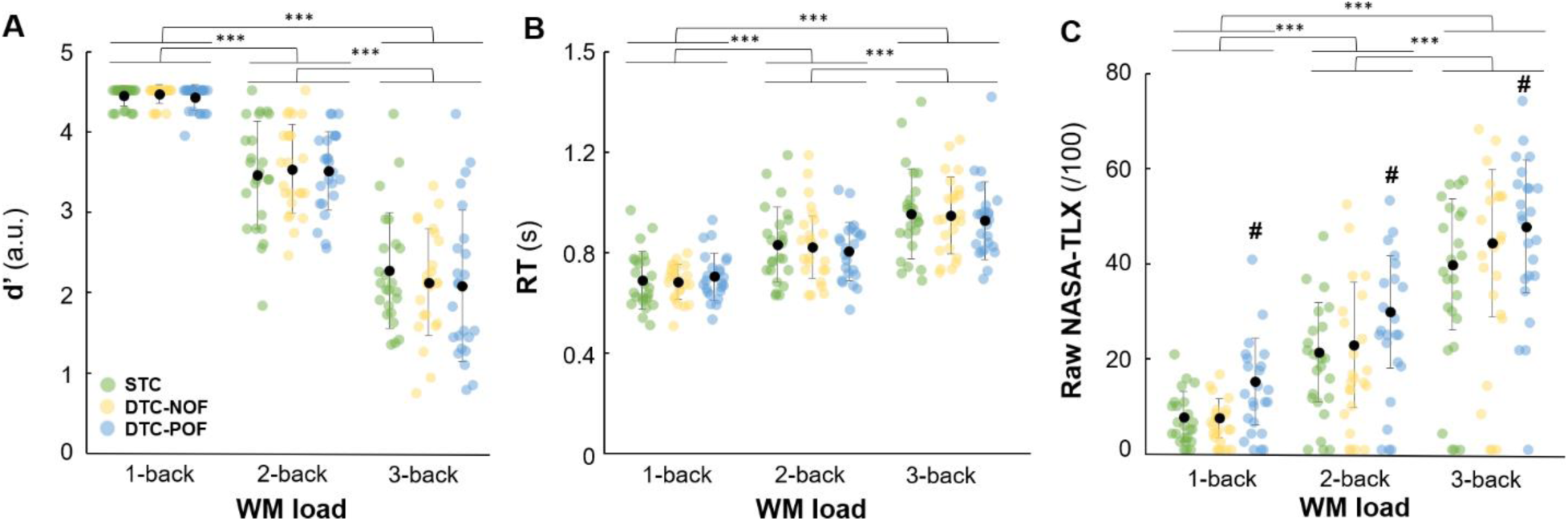
Cognitive performance: A) d-prime (d’, a.u.), B) mean reaction time for target trials (RT, second), and C) raw NASA task load index (Raw NASA-TLX, the maximum score is 100), for the three N-back tasks, *i.e.* 1-back, 2-back, 3-back, performed under three cognitive conditions, *i.e.* single-task cognitive performance in a seated position (STC, green circles), dual-task cognitive (DTC) performance while walking with normal optic flow (DTC-NOF, yellow circles) and perturbed optic flow (DTC-POF, blue circles). Each dot represents a participant, while the black dots and error bars correspond to the population means and standard deviations, respectively. ***: significant differences (p < .001) between the three task loads. #: significant differences (p < .001) between DTC-POF and the other conditions (STC and DTC-NOF).

### Gait kinematics

The mean and standard deviation of all dependent gait kinematic variables for each condition are reported in Table S2.

#### Gait steadiness (mean)

A significant main effect of *Optic flow* was found for mean step width (Table 2, *Tables S2 and S3 for descriptive data*). Post hoc tests revealed an increase in mean step width in POF conditions compared with NOF conditions. These results indicated that participants widened their steps to enhance gait balance. No interaction effect of *Optic flow* x *Walking condition* was observed, indicating that the addition of POF did not influence gait steadiness, regardless of whether walking was performed under single-task or dual-task conditions, and irrespective of concomitant WM load (Figure S2A).

**Table 2.**
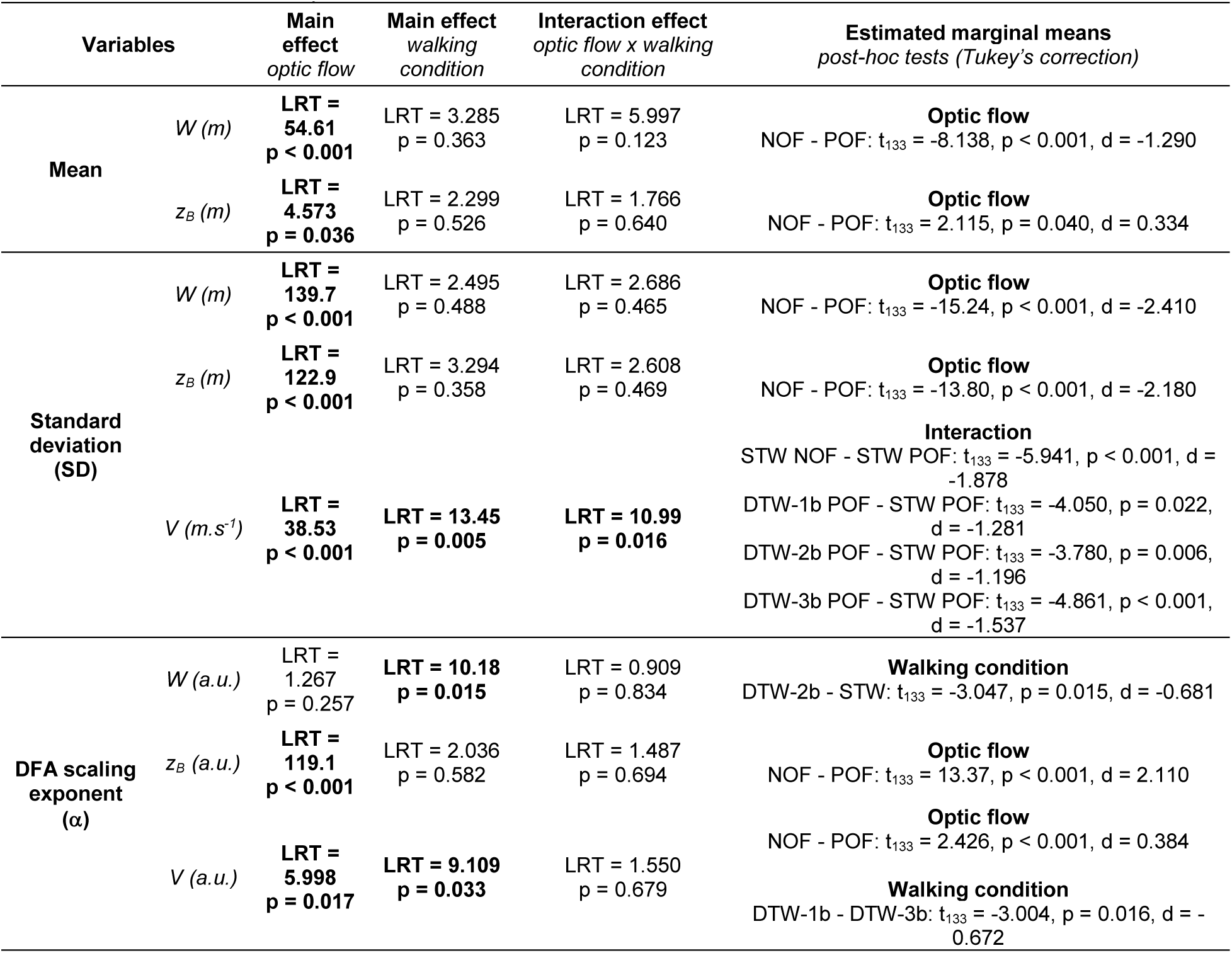
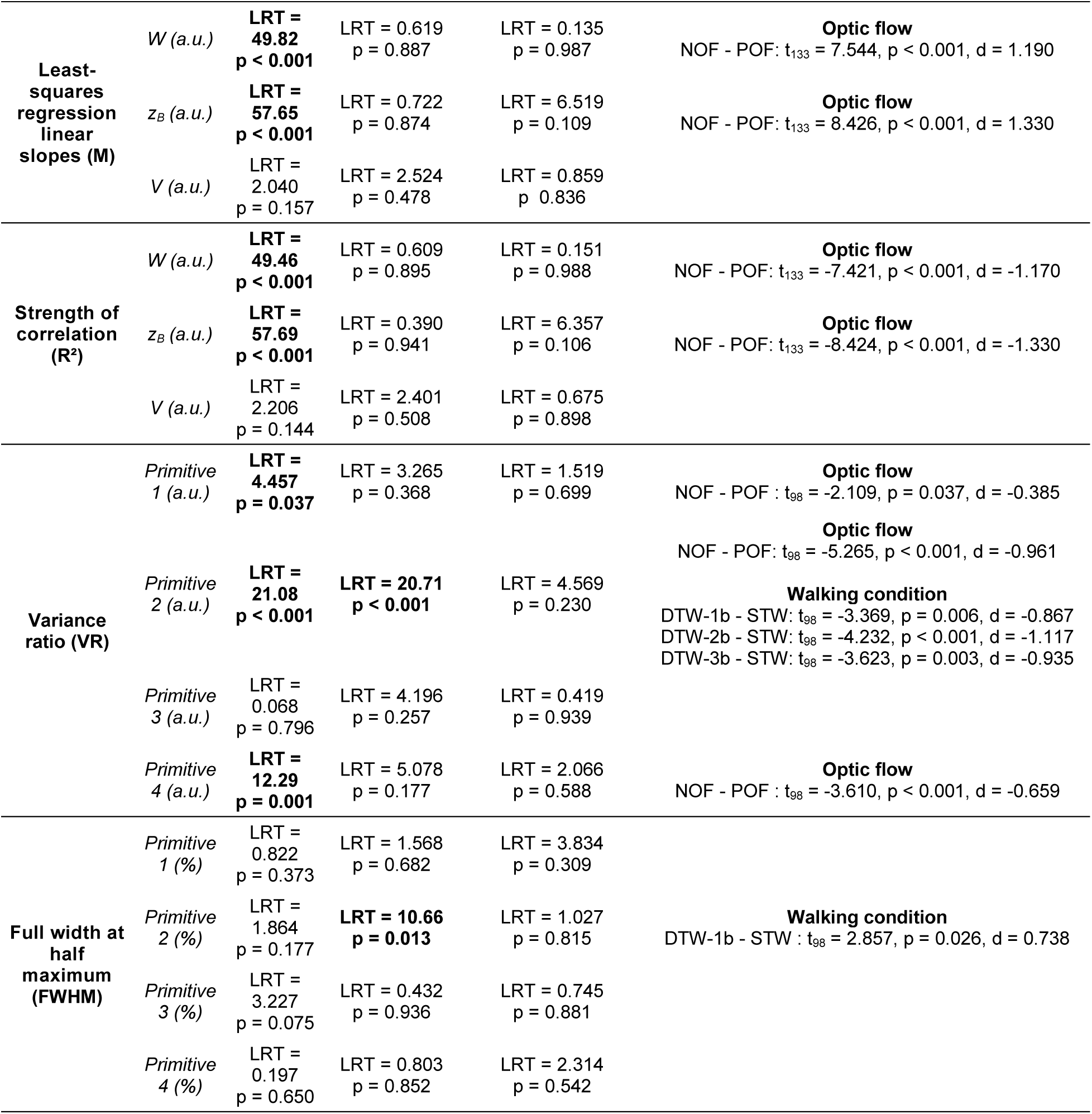
Statistical results for gait dependent variables: kinematics, *i.e.* steadiness (mean), variability (SD: standard deviation) and complexity (α: alpha exponent, M: least-squares regression linear slope, R²: strength of correlation) of step width (W), lateral body position (z _B_) and step velocity (V), and electromyography, *i.e.* motor primitive variability (VR: variance ratio) and motor primitive duration (FWHM: full width at half maximum).

#### Gait variability (standard deviation)

A significant main effect of *Optic flow* was found for standard deviation of step width, lateral body position and step velocity (Table 2, Figure S2A). Post hoc tests revealed an increase in standard deviation values (*i.e.* variability) in POF conditions compared with NOF conditions, indicating that POF induced greater gait variability (Figure 4).

**Figure 4.**
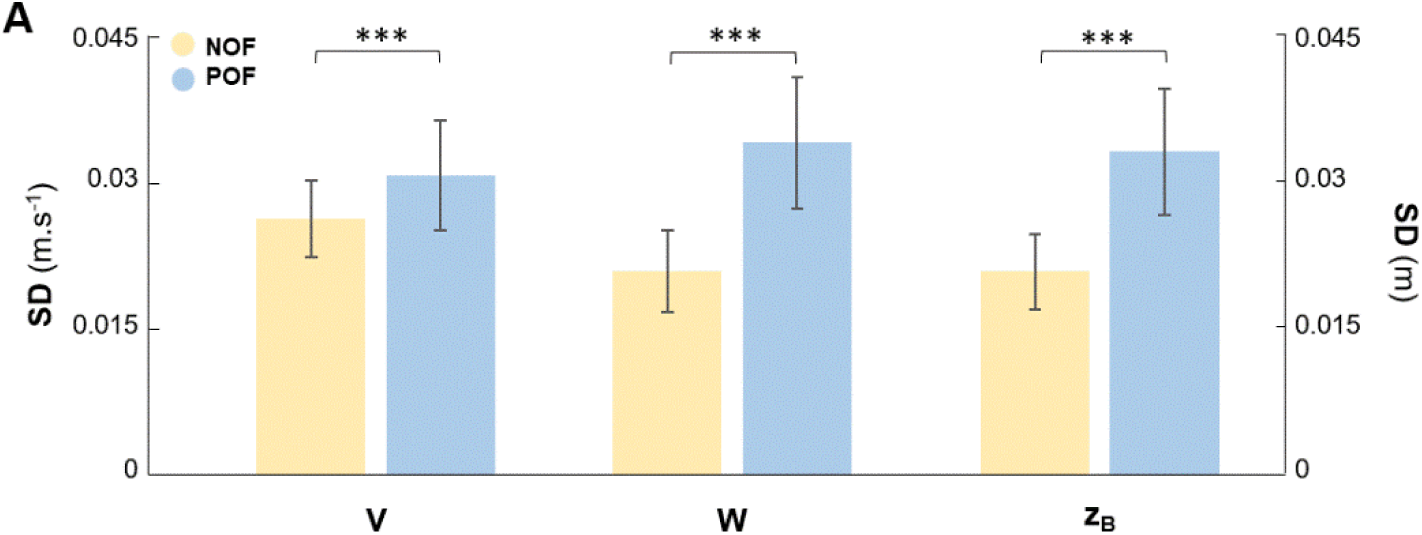
Main effect of *Optic flow* for A) standard deviation (SD) of step velocity (V), step width (W) and lateral body position (z_B_). The two optic flow conditions are represented, i.e. NOF: normal optic flow (yellow bars) and POF: perturbed optic flow (blue bars). ***: significant differences (p < .001) between the two optic flow conditions.

A significant interaction of effect of *Optic flow* x *Walking condition* was found for step velocity. Post hoc tests revealed a reduction in the standard deviation of step velocity only in all POF dual-task walking conditions compared to the POF single-task walking condition, indicating that the addition of a concomitant cognitive task, irrespective of the associated WM load, reduced the impact of POF on gait variability (Figure 5).

**Figure 5.**
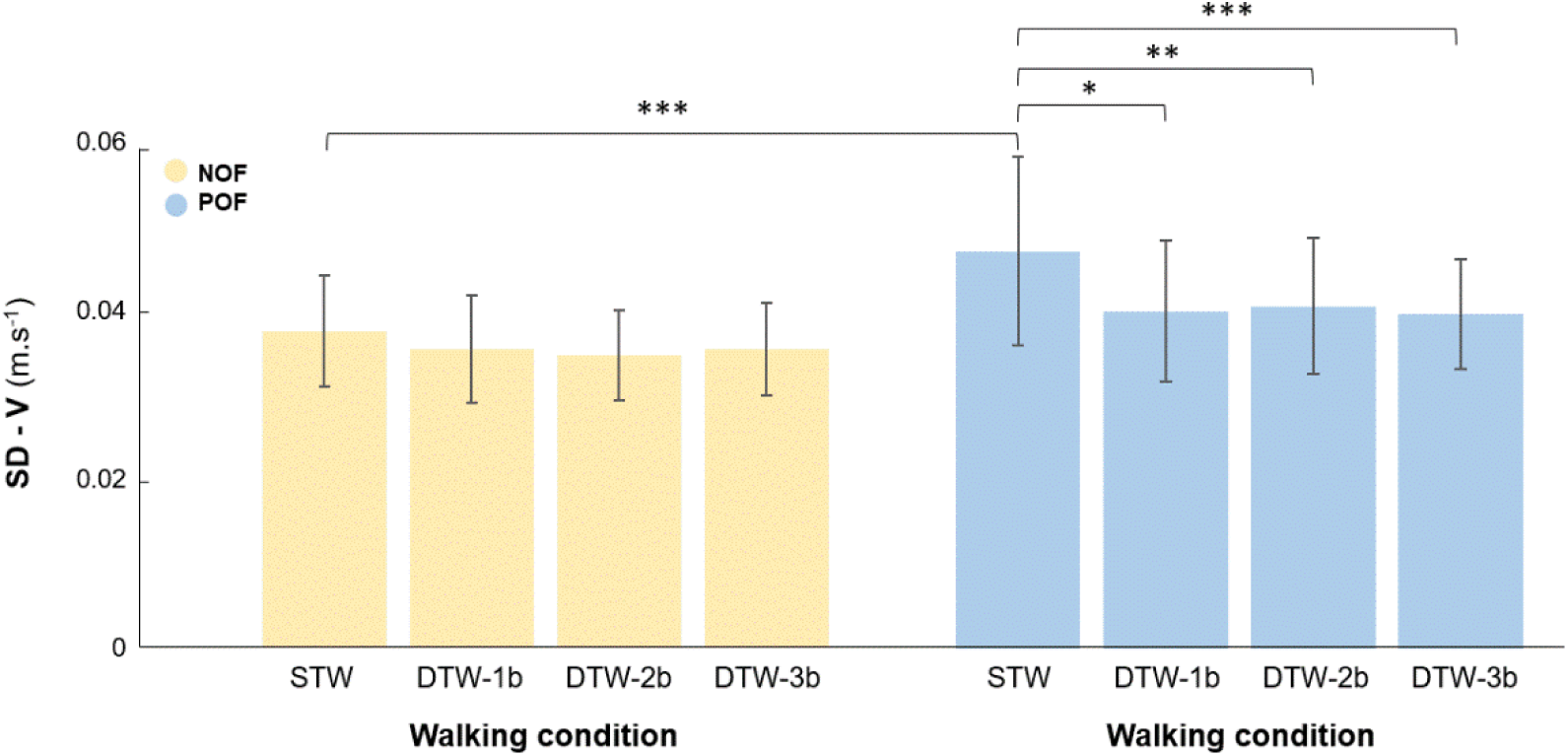
Interaction effect Optic flow x Walking condition for variability (SD: standard deviation) of step velocity (V). The four walking conditions: single-task walking (STW) and dual-task walking (DTW), i.e. walking while simultaneously responding to auditory 1-back (DTW-1b), 2-back (DTW-2b), and 3-back (DTW-3b) tasks, performed under two optic flow conditions, i.e. normal optic flow (NOF, yellow bars) and perturbed optic flow (POF, blue bars) are represented. Significant differences (* p < .05, ** p < .01, *** p < .001) between conditions.

#### Gait complexity (α: alpha exponent, M: least-squares regression linear slope, R^2^: strength of correlation)

A significant main effect of *Optic flow* was found for scaling exponent (α) of lateral body position and of step velocity, slope of step width and of lateral body position (Table 2, Figure S2). Post-hoc tests revealed greater anti-persistence of step velocity (α << 0.5) and lower persistence (α ≈ 0.5) of lateral body position in POF conditions compared with NOF conditions (Figure 6A). Furthermore, they revealed a more negative slope value (closer to −1) and a higher R^2^ value for step width and lateral body position in POF conditions compared with NOF conditions (Figure 6B).

**Figure 6.**
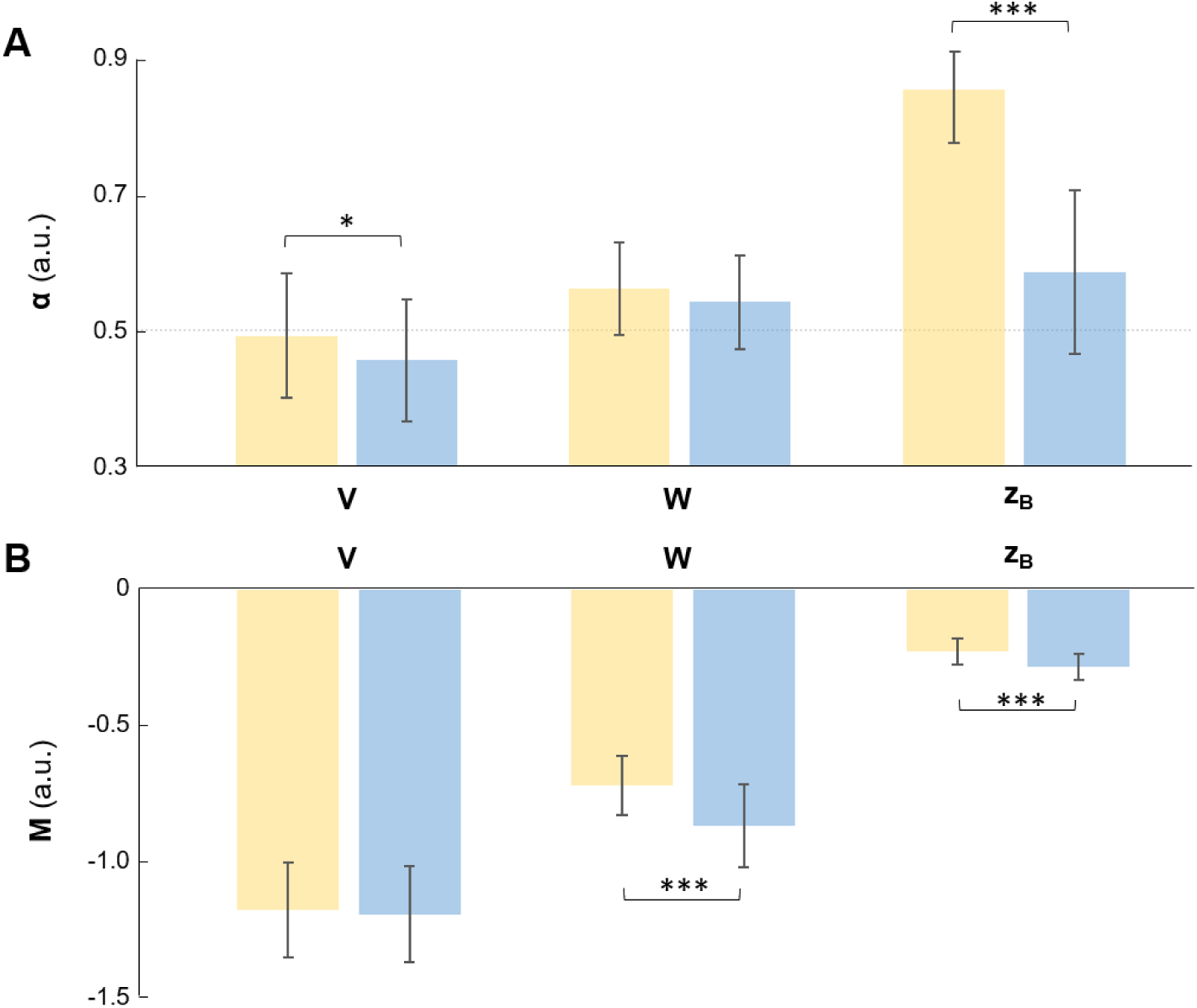
Main effect of *Optic flow* for A) detrended fluctuation analysis scaling exponent (α), B) least-squares regression linear slope (M) of step velocity (V), step width (W) and lateral body position (z_B_). See legend of Figure 4 for details. Significant differences (* p < .05, *** p < .001) between the two optic flow conditions.

These results indicate a tighter control of these variables during perturbed optic flow. A significant main effect of *Walking condition* was found for α of step width and of step velocity (respectively, Figures 7A and 7B). Post hoc tests revealed a decrease in persistence of step width in DTW-2b conditions compared with STW conditions and a decrease in anti-persistence of step velocity (α ≈ 0.5) in DTW-3b conditions compared with DTW-1b conditions. These results indicate that an increase in WM load during walking is first manifested by tighter control of step width, followed by looser control of step velocity.

**Figure 7.**
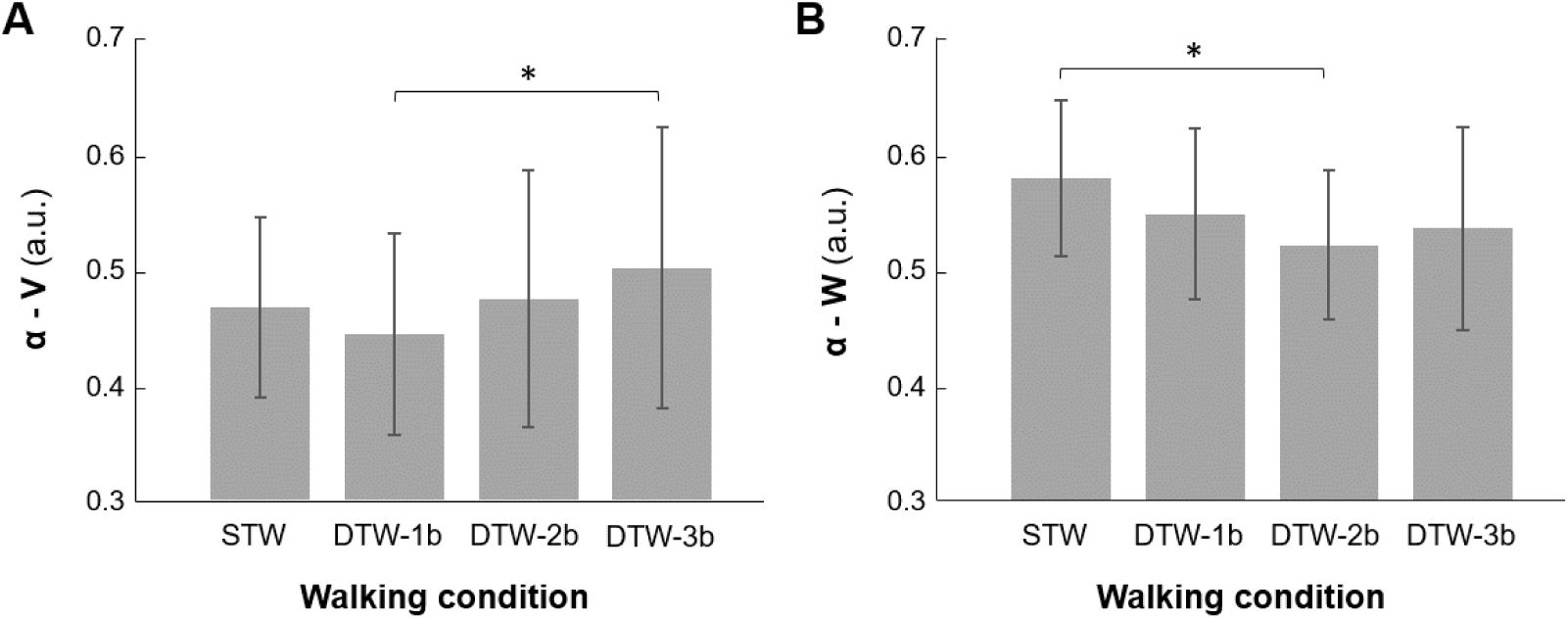
Main effect of *Walking condition* for detrended fluctuation analysis scaling exponent (α) of A) step velocity (V) and B) step width (W). The four walking conditions: single-task walking (STW) and dual-task walking (DTW), *i.e.* walking while simultaneously responding to auditory 1-back (DTW-1b), 2-back (DTW-2b), and 3-back (DTW-3b) tasks are represented regardless of optical flow condition. *: significant differences (p < .05) between the two walking conditions.

### Gait electromyography (EMG)

The mean and standard deviation of all dependent gait EMG variables for each condition are reported in Table S3.

#### Functional organization of muscle synergies

Analysis of the dimensionality of muscle synergies using NNMF revealed that 3 to 4 muscle synergies were sufficient to adequately capture the gait of young adults whatever the experimental condition considered (Figure 8). As the chi² test performed was non-significant (chi² = 1.81; theoretical critical threshold is set at 3.84), the variables were considered independent, and the *Optic Flow* had no significant impact on the number of synergies defined by VAF. Thus, although a model with three synergies was sufficient for several subjects, we chose to retain four synergies (VAF = 90.5 ± 2 %) because this was the most frequent occurrence and it is consistent with the study by Clark et al. (65) which used the same experimental set-up (65).

**Figure 8.**
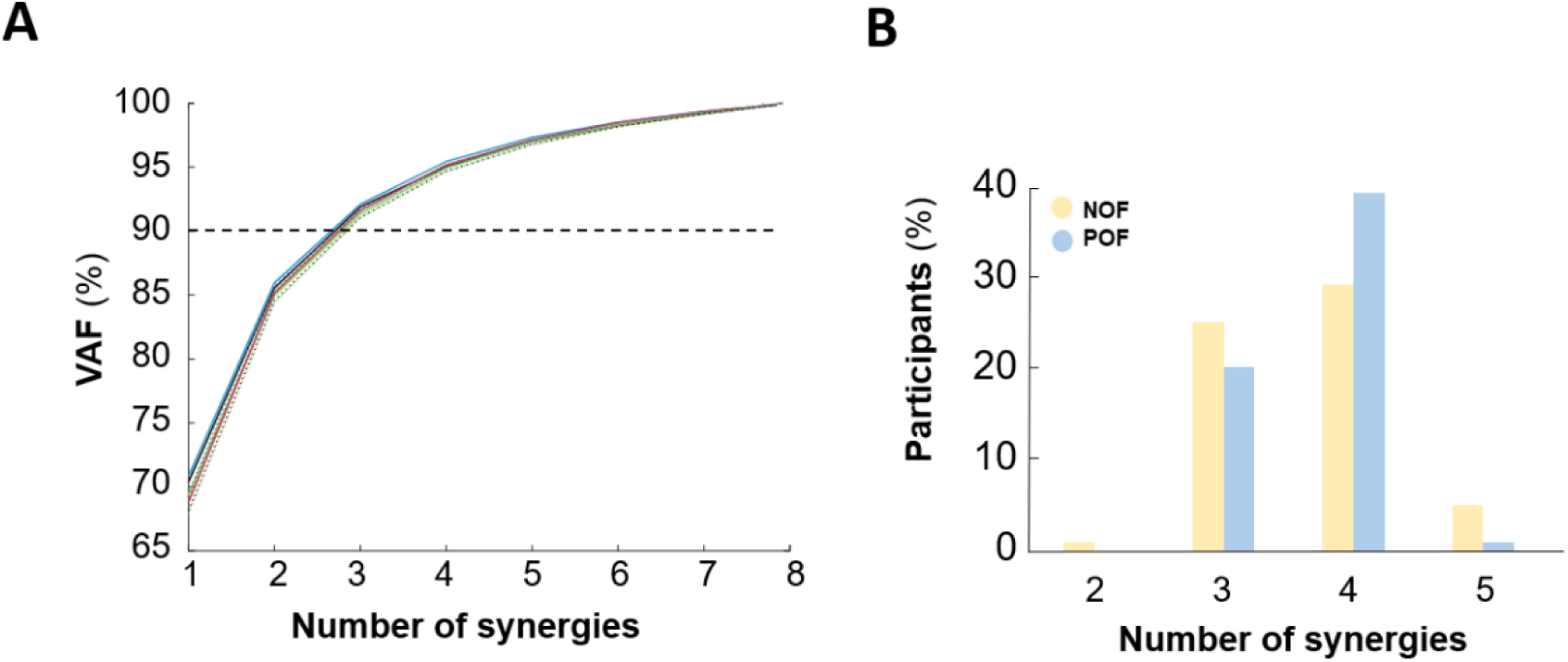
A) Mean variance accounted for (VAF) based on the number of synergies extracted by non-negative matrix factorization (NNMF) for all conditions combined: single-task walking (STW, black) and dual-task walking (DTW) 1-back (DTW-1b, green), 2-back (DTW-2b, blue) and 3-back (DTW-3b, red), performed under normal optic flow (NOF, solid lines) and perturbed optic flow (POF, dotted lines), and B) Percentage of participants (%) as a function of the number of synergies extracted by NNMF under normal optic flow (NOF, yellow bars) and perturbed optic flow (POF, blue bars).

For each synergy, the involvement of each muscle within the synergy (motor module) and its activation profile (motor primitive) are displayed for all conditions, illustrating the relative invariance of both motor modules and motor primitives, whatever the conditions (Figures 9A and 9B). Synergy 1 was active during damping and loading in early stance involving extensor muscles (VM, RF and Gmed). Synergy 2 was composed of the plantar flexor muscles (SOL and GM) and was mainly active during propulsion and swing initiation. Synergy 3 mainly involves the tibialis anterior muscle (TA) and, to a lesser extent, the quadriceps muscles (VM and RF) during the initiation of each phase (stance and swing). Then, Synergy 4 includes activation of the hamstring muscles (BF and ST) in order to ensure the transition between the swing phase and the stance phase by decelerating the leg before foot contact and propelling the body during early stance.

**Figure 9.**
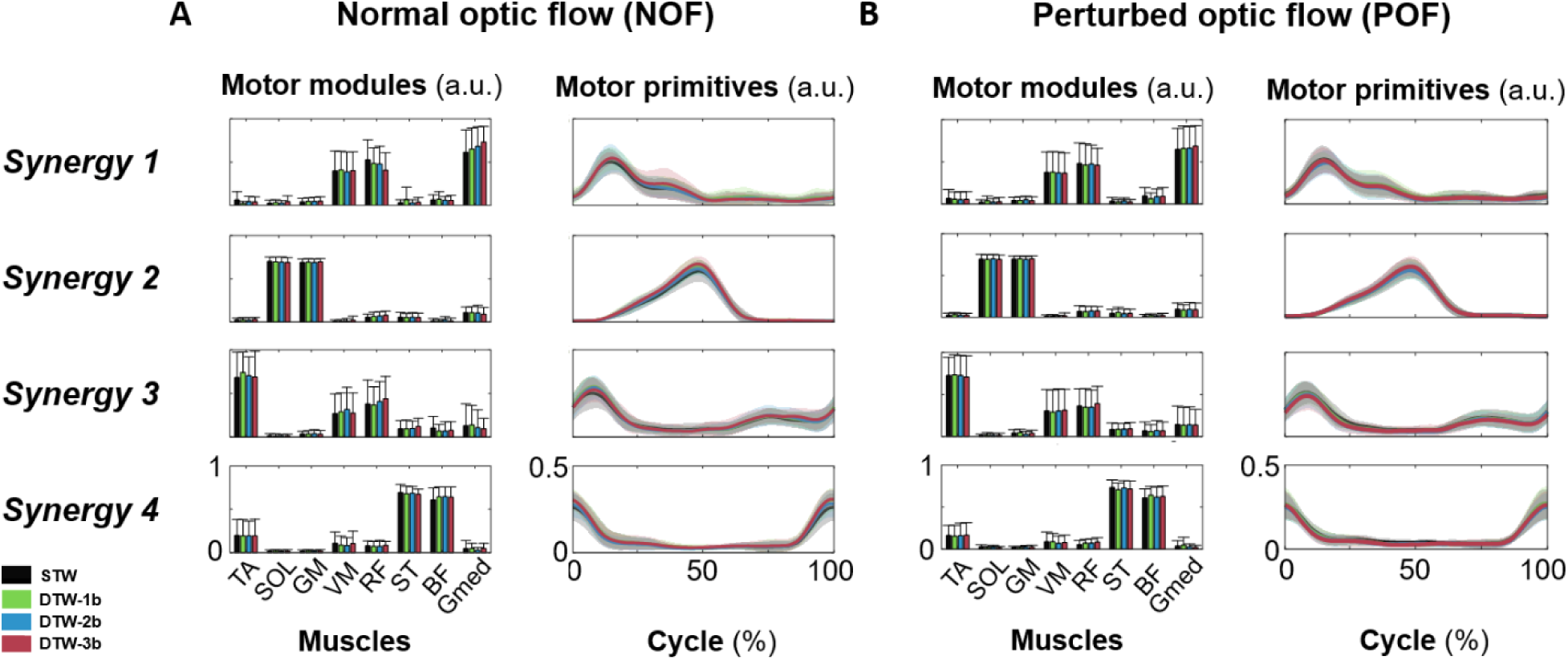
For each synergy, the motor module (spatial component) and motor primitive (temporal component) are displayed for the four walking conditions combined: single-task walking (STW, black) and dual-task walking (DTW) 1-back (DTW-1b, green), 2-back (DTW-2b, blue) and 3-back (DTW-3b, red), performed under A) normal optic flow (NOF) and B) perturbed optic flow (POF).

#### Motor module consistency

A positive correlation was observed for all motor modules for most of the cases, whatever the walking condition. The results did not suggest any impact of WM load (similar modules) on motor module composition (Table 3).

**Table 3.** Motor module similarity (dot product, *r*) observed for each synergy between single-task walking (STW) and each dual-task walking (DTW) conditions, *i.e.* 1-back (DTW-1b), 2-back (DTW-2b) and 3-back (DTW-3b), under normal optic flow (NOF) and perturbed optic flow (POF), respectively. Mean, standard deviation, and percentage of participants for whom *r* > 0.8 are reported.

| Synergy | Variables | NOF |  |  | POF |  |  |
| --- | --- | --- | --- | --- | --- | --- | --- |
|  |  | DTW-1b | DTW-2b | DTW-3b | DTW-1b | DTW-2b | DTW-3b |
| 1 | Mean | 0.92 | 0.88 | 0.86 | 0.88 | 0.92 | 0.98 |
|  | SD | 0.22 | 0.26 | 0.29 | 0.25 | 0.21 | 0.01 |
| | % ($r > 0.8$) | 93% | 87% | 87% | 87% | 93% | 100% |
| 2 | Mean | 0.98 | 0.96 | 0.96 | 0.97 | 0.96 | 0.99 |
|  | SD | 0.06 | 0.09 | 0.09 | 0.07 | 0.08 | 0.01 |
| | % ($r > 0.8$) | 93% | 93% | 93% | 93% | 93% | 100% |
| 3 | Mean | 0.95 | 0.92 | 0.89 | 0.95 | 0.92 | 0.96 |
|  | SD | 0.10 | 0.13 | 0.16 | 0.11 | 0.14 | 0.04 |
| | % ($r > 0.8$) | 93% | 87% | 80% | 93% | 80% | 100% |
| 4 | Mean | 0.93 | 0.93 | 0.95 | 0.95 | 0.94 | 0.96 |
|  | SD | 0.13 | 0.09 | 0.06 | 0.06 | 0.11 | 0.06 |
| | % ($r > 0.8$) | 87% | 93% | 100% | 100% | 93% | 93% |

#### Motor primitive variability (variance ratio) and duration (FWHM: full width at half-maximum)

A significant main effect of *Optic flow* was found for the variance ratio values of motor primitives 1, 2 and 4 (Table 2, Figure S3). Post hoc tests revealed an increase in variance ratio values (*i.e.* variability) in POF conditions compared to NOF conditions (Figure 10).

**Figure 10.**
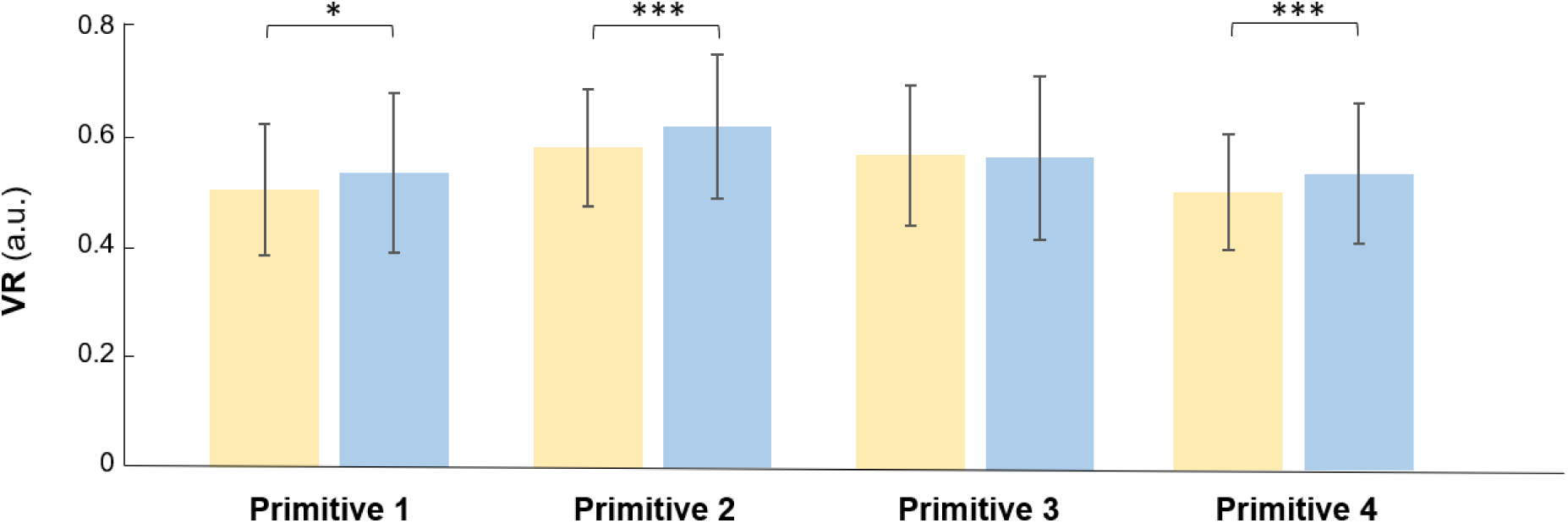
Main effect of *Optic flow* for variance ratio (VR) of each motor primitive. See legend of Figure 4 for details. Significant differences (* p < .05, *** p < .001) between the two optic flow conditions.

A significant main effect of *Walking condition* on variance ratio values was found for motor primitive 2. Post hoc tests revealed a decrease in variance ratio values in DTW-1b, DTW-2b and DTW-3b conditions compared with STW conditions, indicating that participants reduced their variability in muscle synergy activation during propulsion in dual-task conditions (Figure 11A). While there was no effect of *Optic flow* on FWHM values, a significant main effect of *Walking condition* was found for motor primitive 2. Post hoc tests revealed higher FWHM values in DTW-1b conditions compared with STW conditions, indicating that motor primitive duration increased only in low-difficulty dual-task conditions (Figure 11B).

**Figure 11.**
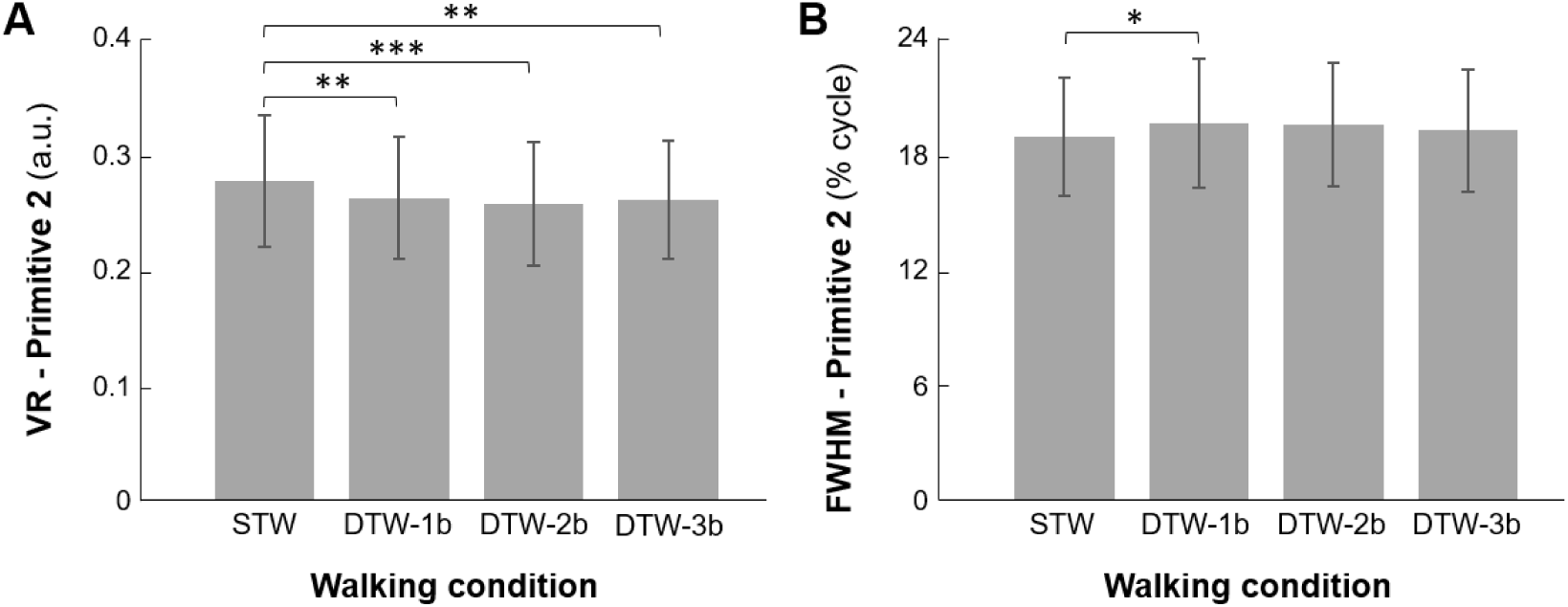
Main effect of *Walking condition* for A) variance ratio (VR, a.u.), and for B) full width at half maximum (FWHM, % cycle) of motor primitive 2. See legend of Figure 7 for details. Significant differences (* p < .05, ** p < .01, *** p < .001) between the two walking conditions.

## DISCUSSION

The aim of this study was to determine cognitive involvement in gait control at kinematic and neuromuscular levels. Overall, the results revealed how young adults adjust (automatic vs. executive control) processes involved in goal-directed locomotion when exposed to varying levels of constraints. Besides, this study identified kinematic and neuromuscular markers that more fully characterize the involvement of executive control in the regulation of task-relevant variables during walking. Specifically, the results confirmed the first hypothesis: kinematic (*i.e.* step parameters) and neuromuscular (*i.e.* motor primitives) variability were greater under POF conditions than in NOF conditions, and young adults sought to counteract perturbations by rapidly correcting task-relevant gait fluctuations. The second hypothesis was partially confirmed: the depletion of cognitive resources through dual-tasking led to a reduction in kinematic and neuromuscular variability and looser regulation of task-relevant gait fluctuations. However, this effect occurred to the same extent regardless of the simultaneous WM load. Furthermore, the duration of the motor primitive of the propulsion-related synergy during the stance phase was found to be increased, but only with the addition of a low-demand secondary task (1-back). Interestingly, and in line with the existing literature, increasing WM load led to a prioritization of gait control in the mediolateral direction over the anteroposterior direction. Finally, the third hypothesis was partially supported, since the impact of POF on kinematic variability (step velocity) was reduced when a cognitive task was performed simultaneously, but this phenomenon was not more pronounced with an increase in WM load, on the one hand, and was not found for other kinematic and neuromuscular variables, on the other. An explanation for this seemingly counterintuitive result is proposed later in the discussion.

As expected, cognitive performance decreased (greater RT and lower d’) and subjective mental workload increased (greater raw NASA-TLX score) with increasing WM load as previously found (52, 74, 75). These results validated our incremental dual-task paradigm (3-back > 2-back > 1-back) under both single-task and dual-task conditions. In addition, participants reported a higher subjective mental workload in POF condition than in NOF and sitting (single-task) conditions. However, as Pechtl et al. (27), cognitive performance did not differ between the sitting and walking conditions, indicating that participants maintained their cognitive engagement and performance regardless of the context. Conversely, this result contrasts with those of Kao and Pierro (41) who observed dual-task effects on cognitive performance during perturbed walking (*i.e.* continuous treadmill platform sways). The lack of interference of concurrent WM load on cognitive performance could be due to the perturbation modality, visual *vs.* mechanical, which involves different mechanisms, and therefore probably does not require the same demand in terms of attentional resources (25, 76). Hence, it is likely that the complexification of the locomotor task, through the use of visual perturbations, does not represent a sufficiently large threat to healthy young adults, unlike mechanical perturbations, so that they are able to maintain their engagement in the performance of a simultaneous cognitive task. Nevertheless, the fact that performance on the concomitant cognitive task was not altered makes it easier to interpret the adjustments made to gait control.

The effects of POF on gait control were observed at the kinematic and neuromuscular levels. In line with previous studies, variability in mediolateral (lateral body position, step width) and anteroposterior (step velocity) kinematic parameters increased under POF (25, 77–80). This increased variability was also demonstrated at the neuromuscular level with the increased variance ratio for three (1,2 and 4) of the four muscle synergies. To our knowledge, this is the first demonstration in the literature of a joint increase in variability at the kinematic and neuromuscular levels following exposure of the locomotor system to a perturbation.

To overcome gait variability induced by POF, young adults exert greater executive control in order to preserve gait balance (26). This is observed through an increase of step width (*i.e.* a wider base of sustentation; 41, 79), tighter regulation (*i.e.* an increase in step-by-step corrections) of lateral body position and step width, and a reduction in persistence of fluctuations in lateral body position (25, 26) under POF conditions. Furthermore, as in previous studies, corrections were tighter for step width than for lateral body position in all conditions (23, 25). Overall, these findings are consistent with multi-objective control simulations (23), according to which young adults increase CNS control by further correcting deviations in step width and lateral body position in order to maintain a central position on the treadmill. Lastly, the antipersistence of step velocity fluctuations was increased in POF conditions (α < 0.5), indicating tighter regulation of these fluctuations. These observations suggest an overcorrection typical of less optimal regulation (9, 60, 82). Taken together, these results provide further evidence that cognitive involvement in gait control increases in the presence of a perturbation, here in the visual modality.

Nevertheless, contrary to expectations, neither the number of motor modules nor the duration (FWHM) of motor primitives changed under POF; hence, the CNS maintains gait control by recruiting a limited number of robust motor synergies. However, the activation profiles of these synergies were more variable in response to POF. According to Desrochers et al. (17), the full expression and proper timing of muscle synergies for mammalian locomotion requires inputs from supraspinal structures and/or limb afferents. Thus, one hypothesis is that, in humans, the greater variability in activation profiles of synergies could potentially reflect POF-related disturbances in these inputs.

The interference effect of a concurrent WM task on gait control was observed at both kinematic and neuromuscular levels. Precisely, at the neuromuscular level, an increase in the duration of motor primitive 2 *(i.e.* plantar flexors) and a decrease in its variability were observed from the first level of WM load (*i.e.* DTW-1b, which mainly involves external focus of attention and relatively few central executive resources). These results suggest that participants stiffened their gait control by modifying propulsion-related synergy during the stance phase when faced with a simultaneous WM task, presumably to make treadmill walking more stable and safer, as previously suggested (16). This more rigid gait pattern under dual-task conditions proved effective, as evidenced by the absence of changes in gait steadiness (*i.e.* mean values of kinematic parameters). In summary, only plantar flexor activity during the propulsive phase was modulated by the CNS in response to increased WM load. These results highlight the CNS’s flexibility to modulate a single synergy without affecting others (*i.e.* synergies 1, 3 and 4), as previously observed in walk-run transition tasks (69, 83). Interestingly, the increase in the duration of motor primitive 2 was only observed in DTW-1b, not in DTW-2b and DTW-3b, with the latter two inducing greater competition between the locomotor and WM tasks at hand for limited attentional resources. These results corroborate those of Walsh (18), who found no effect of a backward counting task (which is also an executive task like the 2 - and 3-back tasks) on FWHM values (note that FWHM values were calculated across all synergies, unlike in our study). A likely explanation is that, under DTW-1b condition, parallel task processing is possible because the cognitive task requires relatively few central executive resources. Thus, the CNS was able to allocate some of these resources to motor control adjustments to increase gait balance. Conversely, under 2 - and 3-back conditions, it is probable that most of the central executive resources were allocated to the maintenance of cognitive performance, leaving insufficient resources for such adjustments (80, 81).

This proposition is partially confirmed by the kinematic data. With the concomitant increase in WM load during walking, fluctuations in step width became more tightly regulated from STW condition to DTW-2b condition. This tighter regulation was certainly enabled through the involvement of central executive processes (9, 82). Taken together, the decreased persistence of step width fluctuations and the increased duration of the motor primitive 2 involving the plantar flexors are consistent with the adoption of a more rigid gait pattern in the mediolateral direction. Consequently, the search for greater gait balance comes at the cost of lower flexibility. Conversely, fluctuations in step velocity became less tightly regulated from DTW-1b condition to DTW-3b condition; one hypothesis is that participants no longer had sufficient central executive resources to correct fluctuations in step velocity in the anteroposterior direction as rapidly. This phenomenon was also observed in a previous study where WM load was manipulated using a dichotic listening paradigm (9). Collectively, these findings provide further evidence that central executive processes are involved in regulating gait fluctuations relevant to the locomotor task goal (*i.e.* maintaining constant step velocity and lateral body position in the center of the treadmill) and that, with increasing WM load, participants prioritize gait control in the mediolateral direction (*i.e.* tighter control) at the expense of control in the anteroposterior direction (*i.e.* looser control) which is neglected. Such a strategy allows for more efficient use of central executive resources, based on the assumption that gait possesses passive dynamic stability in the anteroposterior direction and that active control is critical for maintaining balance in the mediolateral direction (84, 85).

Lastly, an interaction effect was only observed for step velocity variability. This raises the question of why such an interaction effect was found for this particular variable. A likely explanation is that the visual constraint (*i.e.* POF) applied in the mediolateral direction increased step velocity variability by interfering with the mechanical constraint (*i.e.* treadmill moving at a constant speed) applied in the anteroposterior direction. Young adults, by directing their attentional resources to the concurrent auditory task, may have paid less attention to visual information from the virtual environment, thus mitigating the impact of POF on gait control. Besides, from a methodological point of view, the use of a conservative statistical method (*i.e.* linear mixed-effects model with bootstrapping) may have limited interaction effects in our analyses compared to other studies (4, 25, 27).

A few methodological issues should be considered when interpreting the results of this study. First, STW conditions were systematically performed before DTW ones; indeed, STW conditions performed after the DTW ones were not included in the main analyses for the reasons mentioned in the *Supplementary material S1*. However, substantial initial familiarization with treadmill walking was provided to limit any possible learning effect. Second, treadmill walking was performed at a fixed speed (although self-selected by the participant). The differences between fixed-speed treadmill walking and overground walking (*i.e.* everyday walking) have been extensively studied and highlighted in many studies (86, 87). For example, some studies have observed a decrease in variability of several kinematic parameters during treadmill walking compared to overground walking (88, 89). This phenomenon has not prevented the observation of the impact of visual and cognitive constraints on gait variability and, more broadly, on gait regulation. Furthermore, the use of fixed-speed treadmill walking was intended to explore the impact of experimental constraints on variables known to be regulated by the locomotor system, *i.e.* variables that allow satisfying the (implicit) goal of the locomotor task (“not to go beyond the anteroposterior and mediolateral limits of the treadmill”). Finally, we did not include the adductors in our analysis. Given our hypothesis and experimental setup, the motor primitive duration would certainly have been affected. Nevertheless, we defined our muscle set based on the literature to ensure comparability with previous studies. Furthermore, although the muscles selected in the present study are primarily involved in anteroposterior control, some also function as rotators and may be involved in mediolateral control.

In conclusion, this study replicated and extended previous research on how humans regulate gait from step to step to achieve goal-directed walking (23, 60, 62), through the use of a virtual reality-induced optic flow disruption paradigm to destabilize lateral walking balance and a dual-task paradigm to progressively deplete attentional resources that can be allocated to walking. These two paradigms have been used separately and together to gain a deeper understanding of how cognitive processes intervene in gait (stepping) regulation. This study has several strengths. To our knowledge, it is the first study to more comprehensively characterize cognitive contributions to walking by focusing on potentially relevant gait variables and assessing various aspects (*i.e.* steadiness, variability, complexity), and linking kinematic responses to neuromuscular responses. The results revealed that humans optimally regulate their gait to effectively cope with interacting task and environmental constraints, by maintaining relatively stable muscle synergies, flexibly allocating attentional resources between the two tasks at hand, and prioritizing gait variables most relevant to the specific goal-directed locomotor tasks. Specifically, an increase in WM load during walking led to an initial increase in the motor primitive duration of propulsion-specific synergy, followed by stiffer control in the mediolateral direction and looser control in the anteroposterior direction. These findings encourage future research to examine compensatory strategies employed by older adults or by patients with peripheral or central deficits, to counteract the negative effects of aging or underlying pathological processes on gait control in unstable environments.

## DATA AVAILABILITY

All data collected are available and can be downloaded from Dryad platform (http://doi.org/10.5061/dryad.bnzs7h4ds).

## ACKNOWLEDGMENTS

We are very grateful to the participants who took part in the study, and to the Interdisciplinary Center for Virtual Reality (CIREVE) in Caen for their unwavering help and support.

## AUTHOR CONTRIBUTIONS

L.M.D., J.F., V.L. designed the whole study; N.L. conceived the virtual environment; L.M.D., J.F., V.L. conceived the data analysis plan; V.C., V.L., T.R. collected the data; V.C., V.L. analyzed the data; L.M.D., J.F., V.L. interpreted the results; L.M.D., J.F., V.L. wrote the manuscript. L.M.D., J.F., V.L., E.V. obtained funding. All authors read and approved the final version of the manuscript before submission.

