## Supplementary materials for "Kinematic and neuromuscular characterization of cognitive involvement in gait control in healthy young adults"

**Supplementary Material**

An *Optic flow* (NOP, POF) x *Attempt* (1, 2) linear mixed-model (with random intercept for each participant) showed a significant main effect of *Optic flow* (LRT = 48.83; $p < 0.001$), *Attempt* (LRT = 8.049; $p = 0.006$) and interaction effect (LRT = 9.487; $p = 0.002$) on the raw NASA-TLX score. Post-hoc tests on estimated marginal means revealed an increase in subjective mental workload under POF compared to NOF condition ($t_{68.2} = -8.755$, $p < 0.001$, $d = -1.800$) and a decrease in the second attempt compared to the first attempt ($t_{68.2} = 4.190$, $p < 0.001$, $d = 0.863$). More particularly, the raw NASA-TLX score was reduced between attempts 1 and 2 under POF condition ($t_{68.3} = 5.136$, $p < 0.001$, $d = 1.503$) while it was not under NOF condition ($t_{68.2} = 0.774$, $p = 0.866$, $d = 0.223$). The subjective mental workload was almost null under NOF condition, thus the main effects of *Optic flow* and *Attempt* were primarily observed under POF condition. An explanation could be that either a decrease in vigilance (i.e., cognitive relaxation; 1, 2) happened at the end of the blocks or a learning effect (treadmill acclimatization; 3) occurred. Because these effects interfere with the primary objective of the experiment, the second STW was not included in the analysis.

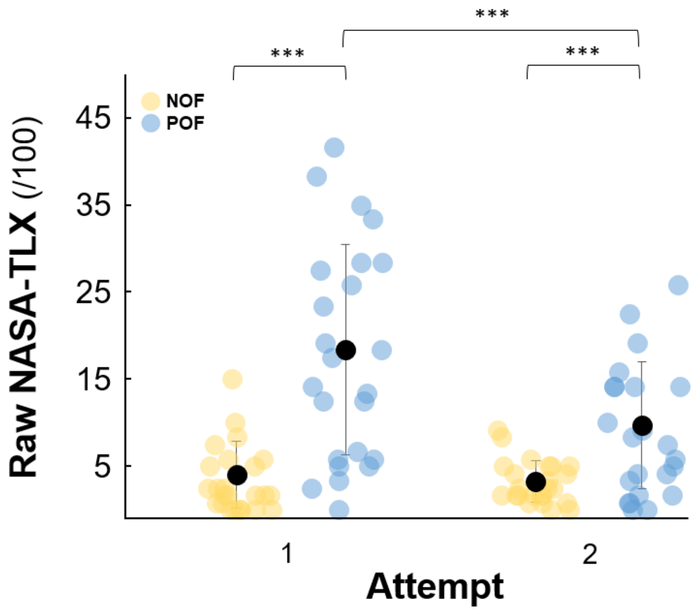


**Figure S1**. Raw NASA-TLX score obtained in two attempts of dual-task walking (1, 2) under normal optic flow (NOF, yellow circles) and perturbed optic flow (POF, blue circles). *** $p < .001$.

**Table S1**. Mean ± standard deviation of cognitive dependent variables for the three cognitive conditions: single-task cognitive performance (STC), dual-task cognitive performance under normal optic flow (DTC-NOF) and dual-task cognitive performance under perturbed optic flow (DTC-POF) and three working memory loads: 1-back (1b), 2-back (2b), and 3-back (3b). Cognitive dependent variables were cognitive task performance, *i.e.* mean reaction time for target trials (RT, second) and d-prime (d', a.u.), and subjective mental workload, *i.e.* raw NASA-TLX score.

| **Variables** | **STC** | | | **DTC-NOF** | | | **DTC-POF** | | |
|---|---|---|---|---|---|---|---|---|---|
| | **1b** | **2b** | **3b** | **1b** | **2b** | **3b** | **1b** | **2b** | **3b** |
| **d'** (a.u.) | 4.451 ± 0.126 | 3.465 ± 0.672 | 2.273 ± 0.719 | 4.473 ± 0.112 | 3.54 ± 0.557 | 2.128 ± 0.664 | 4.428 ± 0.16 | 3.515 ± 0.493 | 2.088 ± 0.941 |
| **RT** (s) | 0.69 ± 0.114 | 0.831 ± 0.151 | 0.955 ± 0.179 | 0.682 ± 0.07 | 0.822 ± 0.125 | 0.948 ± 0.153 | 0.705 ± 0.094 | 0.805 ± 0.116 | 0.927 ± 0.154 |
| **Raw NASA-TLX** (%) | 6.818 ± 5.237 | 20.397 ± 10.48 | 38.889 ± 13.797 | 6.583 ± 4.091 | 21.875 ± 13.228 | 43.333 ± 15.463 | 14.246 ± 9.117 | 28.968 ± 11.826 | 46.884 ± 13.918 |

**Table S2**. Mean ± standard deviation of gait dependent variables for the four walking conditions: single-task walking (STW) and dual-task walking (DTW), *i.e.* walking while simultaneously responding to auditory 1-back (DTW-1b), 2-back (DTW-2b), and 3-back (DTW-3b) tasks, performed under normal optic flow (NOF). Gait dependent variables were kinematics, *i.e.* steadiness (mean), variability (SD: standard deviation) and complexity ($\alpha$: alpha exponent, M: least-squares regression linear slopes, $R^2$: strength of correlation) of step width (W), lateral body position ($z_B$) and step velocity (V), and electromyography, *i.e.* motor primitive variability (VR: variance ratio, a.u.) and motor primitive duration (FWHM: full width at half maximum, % cycle).

| Variables | | NOF | | | |
|---|---|---|---|---|---|
| | | **STW** | **DTW-1b** | **DTW-2b** | **DTW-3b** |
| **Step width (W)** | *Mean (m)* | 0.099 ± 0.028 | 0.100 ± 0.028 | 0.103 ± 0.029 | 0.102 ± 0.028 |
| | *SD (m)* | 0.024 ± 0.005 | 0.024 ± 0.005 | 0.023 ± 0.005 | 0.023 ± 0.004 |
| | *α (a.u.)* | 0.594 ± 0.082 | 0.569 ± 0.103 | 0.531 ± 0.110 | 0.534 ± 0.104 |
| | *M (a.u.)* | -0.725 ± 0.109 | -0.722 ± 0.125 | -0.730 ± 0.143 | -0.741 ± 0.150 |
| | *$R^2$ (a.u.)* | 0.362 ± 0.054 | 0.362 ± 0.062 | 0.365 ± 0.071 | 0.371 ± 0.076 |
| **Lateral body position ($z_B$)** | *Mean (m)* | -0.020 ± 0.017 | -0.016 ± 0.021 | -0.017 ± 0.021 | -0.017 ± 0.021 |
| | *SD (m)* | 0.021 ± 0.004 | 0.021 ± 0.004 | 0.020 ± 0.004 | 0.022 ± 0.006 |
| | *α (a.u.)* | 0.853 ± 0.086 | 0.871 ± 0.143 | 0.815 ± 0.114 | 0.843 ± 0.138 |
| | *M (a.u.)* | -0.227 ± 0.056 | -0.238 ± 0.068 | -0.239 ± 0.059 | -0.221 ± 0.065 |
| | *$R^2$ (a.u.)* | 0.113 ± 0.028 | 0.119 ± 0.034 | 0.118 ± 0.029 | 0.110 ± 0.031 |
| **Step velocity (V)** | *Mean ($m.s^{-1}$)* | 1.117 ± 0.132 | 1.117 ± 0.132 | 1.117 ± 0.132 | 1.117 ± 0.132 |
| | *SD ($m.s^{-1}$)* | 0.037 ± 0.006 | 0.035 ± 0.006 | 0.034 ± 0.005 | 0.035 ± 0.005 |
| | *α (a.u.)* | 0.481 ± 0.094 | 0.450 ± 0.114 | 0.496 ± 0.130 | 0.531 ± 0.139 |
| | *M (a.u.)* | -1.210 ± 0.228 | -1.214 ± 0.201 | -1.192 ± 0.220 | -1.154 ± 0.186 |
| | *$R^2$ (a.u.)* | 0.606 ± 0.114 | 0.607 ± 0.100 | 0.596 ± 0.109 | 0.578 ± 0.092 |
| **Variance ratio (VR)** | *Primitive 1 (a.u.)* | 0.520 ± 0.169 | 0.532 ± 0.158 | 0.493 ± 0.144 | 0.483 ± 0.12 |
| | *Primitive 2 (a.u.)* | 0.264 ± 0.049 | 0.255 ± 0.052 | 0.248 ± 0.045 | 0.256 ± 0.047 |
| | *Primitive 3 (a.u.)* | 0.606 ± 0.16 | 0.559 ± 0.131 | 0.559 ± 0.144 | 0.580 ± 0.173 |
| | *Primitive 4 (a.u.)* | 0.522 ± 0.137 | 0.511 ± 0.106 | 0.499 ± 0.106 | 0.500 ± 0.105 |
| **Full width at half maximum (FWHM)** | *Primitive 1 (%)* | 10,493 ± 1,64 | 10,034 ± 1,687 | 10,466 ± 1,63 | 10,508 ± 1,544 |
| | *Primitive 2 (%)* | 18,892 ± 2,836 | 19,399 ± 2,984 | 19,389 ± 3,188 | 18,981 ± 3,143 |
| | *Primitive 3 (%)* | 10,038 ± 1,972 | 10,151 ± 1,819 | 10,647 ± 1,277 | 10,737 ± 1,526 |
| | *Primitive 4 (%)* | 10,769 ± 1,527 | 10,621 ± 1,383 | 10,791 ± 1,632 | 10,708 ± 1,954 |

**Table S3**. Mean ± standard deviation of gait dependent variables for the four walking conditions: single-task walking (STW) and dual-task walking (DTW), *i.e.* walking while simultaneously responding to auditory 1-back (DTW-1b), 2-back (DTW-2b), and 3-back (DTW-3b) tasks, performed under perturbed optic flow (POF). Gait dependent variables were kinematics, *i.e.* steadiness (mean), variability (SD: standard deviation) and complexity ($\alpha$: alpha exponent, M: least-squares regression linear slopes, $R^2$: strength of correlation) of step width (W), lateral body position ($z_B$) and step velocity (V), and electromyography, *i.e.* motor primitive variability (VR: variance ratio, a.u.) and motor primitive duration (FWHM: full width at half maximum, % cycle).

| **Variables** | | **POF** | | | |
|---|---|---|---|---|---|
| | | **STW** | **DTW-1b** | **DTW-2b** | **DTW-3b** |
| **Step width (W)** | *Mean (m)* | 0.113 ± 0.029 | 0.109 ± 0.029 | 0.112 ± 0.030 | 0.110 ± 0.029 |
| | *SD (m)* | 0.041 ± 0.008 | 0.037 ± 0.008 | 0.037 ± 0.009 | 0.038 ± 0.008 |
| | *α (a.u.)* | 0.567 ± 0.108 | 0.529 ± 0.100 | 0.514 ± 0.073 | 0.541 ± 0.096 |
| | *M (a.u.)* | -0.880 ± 0.199 | -0.869 ± 0.165 | -0.873 ± 0.164 | -0.897 ± 0.150 |
| | *$R^2$ (a.u.)* | 0.440 ± 0.099 | 0.434 ± 0.083 | 0.436 ± 0.081 | 0.449 ± 0.075 |
| **Lateral body position ($z_B$)** | *Mean (m)* | -0.020 ± 0.020 | -0.024 ± 0.024 | -0.023 ± 0.024 | -0.020 ± 0.023 |
| | *SD (m)* | 0.036 ± 0.008 | 0.033 ± 0.007 | 0.032 ± 0.007 | 0.033 ± 0.007 |
| | *α (a.u.)* | 0.572 ± 0.136 | 0.624 ± 0.136 | 0.584 ± 0.154 | 0.549 ± 0.159 |
| | *M (a.u.)* | -0.291 ± 0.049 | -0.274 ± 0.050 | -0.291 ± 0.059 | -0.292 ± 0.058 |
| | *$R^2$ (a.u.)* | 0.146 ± 0.025 | 0.136 ± 0.025 | 0.145 ± 0.029 | 0.146 ± 0.029 |
| **Step velocity (V)** | *Mean ($m.s^{-1}$)* | 1.116 ± 0.132 | 1.118 ± 0.132 | 1.117 ± 0.132 | 1.117 ± 0.132 |
| | *SD ($m.s^{-1}$)* | 0.047 ± 0.011 | 0.040 ± 0.008 | 0.040 ± 0.008 | 0.039 ± 0.006 |
| | *α (a.u.)* | 0.453 ± 0.081 | 0.437 ± 0.100 | 0.453 ± 0.137 | 0.472 ± 0.129 |
| | *M (a.u.)* | -1.222 ± 0.156 | -1.195 ± 0.186 | -1.232 ± 0.219 | -1.189 ± 0.207 |
| | *$R^2$ (a.u.)* | 0.611 ± 0.078 | 0.597 ± 0.093 | 0.616 ± 0.109 | 0.594 ± 0.103 |
| **Variance ratio (VR)** | *Primitive 1 (a.u.)* | 0.557 ± 0.156 | 0.534 ± 0.141 | 0.529 ± 0.157 | 0.533 ± 0.156 |
| | *Primitive 2 (a.u.)* | 0.290 ± 0.066 | 0.269 ± 0.054 | 0.267 ± 0.062 | 0.265 ± 0.055 |
| | *Primitive 3 (a.u.)* | 0.600 ± 0.148 | 0.573 ± 0.151 | 0.549 ± 0.143 | 0.563 ± 0.165 |
| | *Primitive 4 (a.u.)* | 0.563 ± 0.146 | 0.525 ± 0.127 | 0.549 ± 0.127 | 0.531 ± 0.134 |
| **Full width at half maximum (FWHM)** | *Primitive 1 (%)* | 10,656 ± 1,55 | 10,634 ± 1,527 | 10,288 ± 1,516 | 10,475 ± 1,341 |
| | *Primitive 2 (%)* | 18,872 ± 3,317 | 19,693 ± 3,73 | 19,614 ± 3,209 | 19,404 ± 3,21 |
| | *Primitive 3 (%)* | 10,46 ± 1,576 | 10,259 ± 1,689 | 10,625 ± 1,349 | 10,53 ± 1,215 |
| | *Primitive 4 (%)* | 10,891 ± 1,764 | 10,98 ± 1,706 | 10,521 ± 1,335 | 10,762 ± 2,03 |

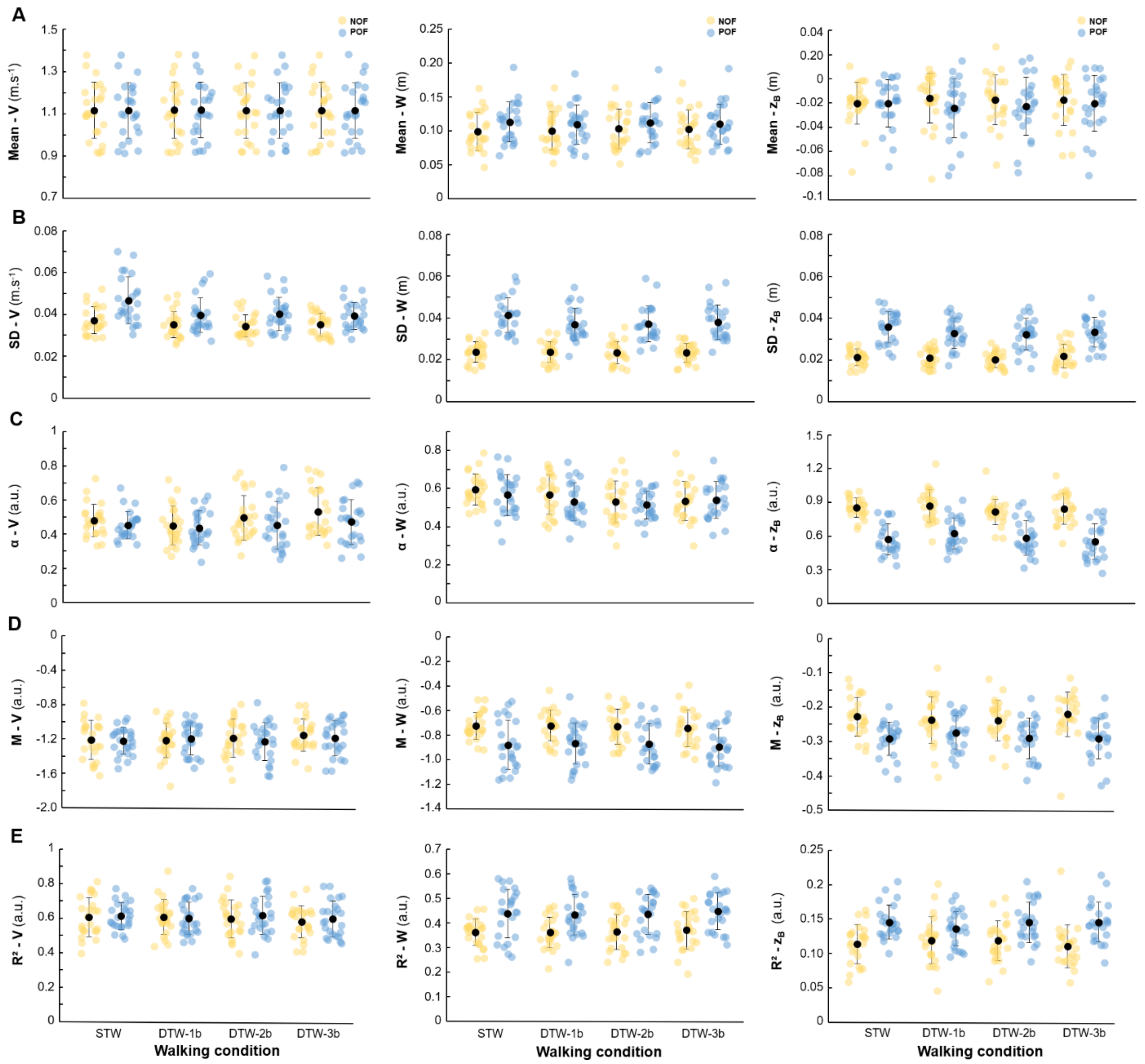


**Figure S2. A)** Mean, **B)** standard deviation (SD), **C)** detrended fluctuation analysis (DFA) scaling exponent (α), **D)** least-squares regression linear slope (M) and **E)** strength of correlation ($R^2$) of step velocity (V, left), step width (W, middle) and lateral body position ($z_B$, right), for the four walking conditions: single-task walking (STW) and dual-task walking (DTW), *i.e.* walking while simultaneously responding to auditory 1-back (DTW-1b), 2-back (DTW-2b), and 3-back (DTW-3b) tasks, performed under two optic flow conditions, *i.e.* normal optic flow (NOF, yellow circles) and perturbed optic flow (POF, blue circles). Each dot represents a participant, while the black dots and error bars correspond to the population means and standard deviations respectively. Significant effects are not represented.

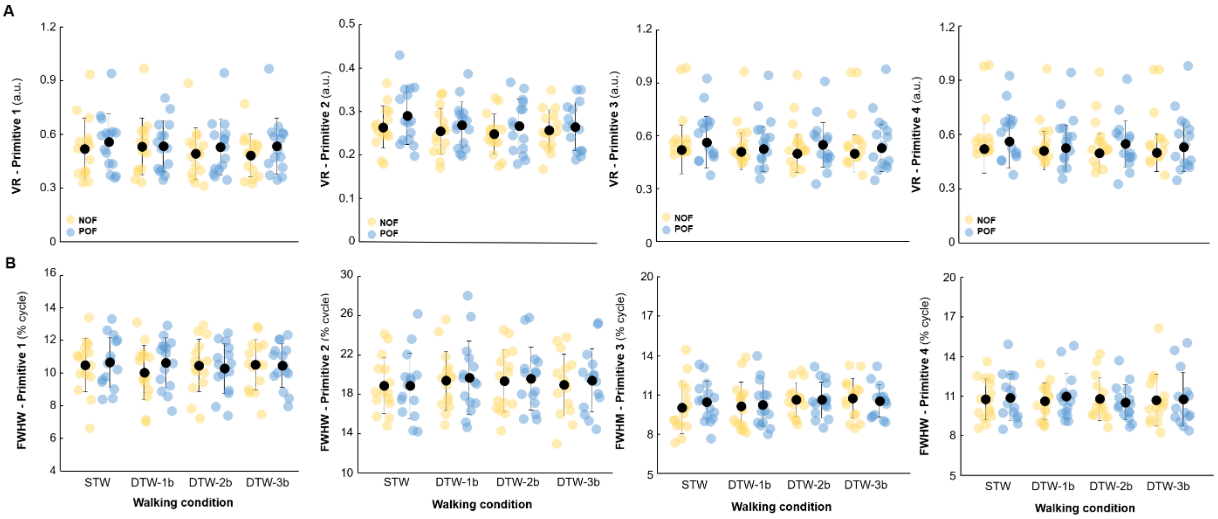


**Figure S3. A)** Variance ratio (VR, a.u.) and **B)** full width at half maximum (FWHM, % cycle) of each motor primitive for the four walking conditions: single-task walking (STW, black) and dual-task walking (DTW) 1-back (DTW-1b), 2-back (DTW-2b) and 3-back (DTW-3b), performed under normal optic flow (NOF, yellow circles) and perturbed optic flow (POF, blue circles). See legend of Figure S2 for details.

### References (additional)


1. **Barwick F**, **Arnett P**, **Slobounov S**. EEG correlates of fatigue during administration of a neuropsychological test battery. *Clin Neurophysiol* 123: 278–284, 2012. doi: 10.1016/j.clinph.2011.06.027.

2. **Benoit C-E**, **Solopchuk O**, **Borragán G**, **Carbonnelle A**, **Van Durme S**, **Zénon A**. Cognitive task avoidance correlates with fatigue-induced performance decrement but not with subjective fatigue. *Neuropsychologia* 123: 30–40, 2019. doi: 10.1016/j.neuropsychologia.2018.06.017.

3. **Meyer C**, **Killeen T**, **Easthope CS**, **Curt A**, **Bolliger M**, **Linnebank M**, **Zörner B**, **Filli L**. Familiarization with treadmill walking: How much is enough? *Sci Rep* 9: 5232, 2019. doi: 10.1038/s41598-019-41721-0.
